# Humoral and cellular immune memory to four COVID-19 vaccines

**DOI:** 10.1101/2022.03.18.484953

**Authors:** Zeli Zhang, Jose Mateus, Camila H. Coelho, Jennifer M. Dan, Carolyn Rydyznski Moderbacher, Rosa Isela Gálvez, Fernanda H. Cortes, Alba Grifoni, Alison Tarke, James Chang, E. Alexandar Escarrega, Christina Kim, Benjamin Goodwin, Nathaniel I. Bloom, April Frazier, Daniela Weiskopf, Alessandro Sette, Shane Crotty

## Abstract

Multiple COVID-19 vaccines, representing diverse vaccine platforms, successfully protect against symptomatic COVID-19 cases and deaths. Head-to-head comparisons of T cell, B cell, and antibody responses to diverse vaccines in humans are likely to be informative for understanding protective immunity against COVID-19, with particular interest in immune memory. Here, SARS-CoV-2-spike—specific immune responses to Moderna mRNA-1273, Pfizer/BioNTech BNT162b2, Janssen Ad26.COV2.S and Novavax NVX-CoV2373 were examined longitudinally for 6 months. 100% of individuals made memory CD4^+^ T cells, with cTfh and CD4-CTL highly represented after mRNA or NVX-CoV2373 vaccination. mRNA vaccines and Ad26.COV2.S induced comparable CD8^+^ T cell frequencies, though memory CD8^+^ T cells were only detectable in 60-67% of subjects at 6 months. Ad26.COV2.S was not the strongest immunogen by any measurement, though the Ad26.COV2.S T cell, B cell, and antibody responses were relatively stable over 6 months. A differentiating feature of Ad26.COV2.S immunization was a high frequency of CXCR3^+^ memory B cells. mRNA vaccinees had substantial declines in neutralizing antibodies, while memory T cells and B cells were comparatively stable over 6 months. These results of these detailed immunological evaluations may also be relevant for vaccine design insights against other pathogens.

## INTRODUCTION

The response to the SARS-CoV-2 pandemic has relied in large part on the development, testing, and deployments of vaccines. In a short time, several different vaccine platforms have been developed and after establishing their safety and efficacy, deployed for use in a large number of individuals. In the USA, two different mRNA vaccines (Moderna mRNA-1273 (Jackson et al., 2020), and Pfizer/BioNTech BNT162b2 (Vogel et al., 2021; Walsh et al., 2020) and a viral vector-based vaccine (Janssen/J&J Ad26.COV2.S) (Sadoff et al., 2021) have been widely used. The recombinant protein-based adjuvanted vaccine Novavax NVX-CoV2373 completed successful Phase 3 efficacy clinical trials in the USA, Mexico, and the UK (Dunkle et al., 2021; Heath et al., 2021) and is approved for use or expected to be approved for use in several different countries (Novavax, 2022). These four vaccines are representatives of the three main vaccine platforms in use for the prevention of COVID-19, namely mRNA, viral vector, and recombinant protein plus adjuvant (Pollard and Bijker, 2021).

In Phase 3 trials, these vaccines proved remarkably effective with early vaccine efficacy (VE) of 95% for BNT162b2 (Thomas et al., 2021), 94% for mRNA-1273 (Baden et al., 2021) and 90% for NVX-CoV2373 (Dunkle *et al*., 2021; Heath *et al*., 2021) against COVID-19 cases. A single dose of Ad26.COV2.S was associated with 67% VE overall and 70% in the USA (Sadoff *et al*., 2021). 6-month efficacy data for BNT162b2 and mRNA-1273 were 91% and 93% against COVID-19 cases (Thomas *et al*.,2021). Population-based “real world” studies of COVID-19 VE have provided additional insights, including comparisons between vaccines. VE wanes against symptomatic COVID-19 over time (Leon et al., 2022; Lin et al., 2022; Pilishvili et al., 2021; Rosenberg et al., 2022; Tartof et al., 2021). In one large study, VE against symptomatic COVID-19 for BNT162b2 and mRNA-1273 decreased to 67% and 75% at 5-7 months (Rosenberg *et al*., 2022). 1-dose Ad26.COV2.S VE started lower and also declined (Rosenberg *et al*., 2022). Comparable findings were made in multiple studies of populations using BNT162b2, mRNA-1273, and Ad26.COV2.S (Leon *et al*., 2022; Lin *et al*., 2022; Rosenberg *et al*., 2022; Tartof *et al*., 2021). If any detectable SARS-CoV-2 infection is considered, as opposed to symptomatic disease, lower VE is observed for all vaccines (Nordstrom et al., 2022; Pouwels et al., 2021). Higher VE against hospitalization is observed for all COVID-19 vaccines, with somewhat lower hospitalization VE for Ad26.COV2.S compared to the mRNA vaccines (e.g. 82% vs. 94% (Rosenberg *et al*., 2022)). Notably, in multiple large “real world” studies, VE against hospitalization was stable over time in contrast to VE against infections (Tartof *et al*., 2021), potentially indicating distinct immunological mechanisms of action contributing to protection against hospitalization compared to detectable infection (Sette and Crotty, 2021).

Antibodies have been established as a clear correlate of protection against infection over the first months post-vaccination (Gilbert et al., 2022; Khoury et al., 2021), but several lines of evidence also suggest important contributions from T and B cell memory responses in protective immunity (Sette and Crotty, 2021), with neutralizing antibodies playing a dominant role in prevention of infection, while cellular immunity might be key to modulate disease severity and resolve infection (Kedzierska, 2022). Overall, available data suggest that coordinated functions of different branches of adaptive immunity may provide multiple mechanisms of protective immunity against COVID-19.

Differences between VE of COVID-19 vaccines suggest that the different vaccines might generate differential immune memory. Comparisons of immunogenicity and immune memory of different COVID-19 vaccines have been limited, hampered by multiple challenges. First, side-by-side comparisons with standardized cellular assays are often lacking. Standardized binding antibody and neutralizing antibody quantitation is possible via the use of WHO international standards (Mattiuzzo, 2020). However, CD4^+^ T cell, CD8^+^ T cell, and memory B cell assays all use live cells and complex reagents, which are far less amenable to cross-laboratory comparisons, and thus memory T and B cell measurements within the same study are required for quantitative comparisons. This is highlighted by the initial discordant findings regarding CD8^+^ T cell responses to COVID-19 mRNA vaccines, with early reports suggesting quite different CD8^+^ T cell response rates to BNT162b2 compared to mRNA-1273 (Corbett et al., 2020; Jackson *et al*., 2020; Sahin et al., 2021). Second, longitudinal studies with cryopreserved PBMCs are needed to directly determine kinetics of vaccine-specific immune memory in humans. Additionally, few studies have assessed antibody, CD4^+^ T cell, CD8^+^ T cell, and memory B cell vaccine responses simultaneously in the same individuals.

The massive COVID-19 immunization campaigns represent a unique opportunity to comprehensively collect and analyze immune responses in a longitudinal fashion for individuals immunized in the same year and having no prior immunity. The present study was designed to establish the magnitude and duration of vaccine-induced immune memory with four different vaccine platforms. A direct, side-by-side, comprehensive evaluation of effector and memory immune responses induced by different vaccine platforms is important to advance our understanding of the protection afforded by the various COVID-19 vaccines, as well as understand fundamental differences in immunogenicity and immune memory to mRNA, adenoviral vector, and recombinant protein vaccine platforms in humans. Here, we compare the immune responses induced by three different vaccine platforms, namely two different mRNA vaccines (Moderna mRNA-1273 and Pfizer/BioNTech BNT162b2), a viral vector-based vaccine (Janssen Ad26.COV2.S) and the protein-based adjuvanted vaccine Novavax NVX-CoV2373. The inclusion of NVX-CoV2373 was of particular interest for head-to-head comparisons of immune memory between a more conventional recombinant protein vaccine and mRNA and viral vectors. We additionally compared their immune memory to natural infection for binding antibodies, neutralizing antibodies, spike-specific CD4^+^ T cells, spike-specific CD8^+^ T cells, and spike- and RBD-specific memory B cells. To the best of our knowledge, this is the most comprehensive side-by-side evaluation of the kinetics of immune memory to these four different vaccine platforms.

## RESULTS

### COVID-19 vaccine cohorts

To compare the development of immune memory, we enrolled subjects who were either planning or had received immunization with mRNA-1273, BNT162b2, Ad26.COV2.S, or NVX-CoV2373 vaccine. Blood donations were obtained at multiple time points, and both plasma and peripheral blood mononuclear cells (PBMC) were preserved. Sampling time points were pre-vaccination (T1), 2 weeks (15 ± 3 days) after the 1^st^ immunization (T2), 45 ± 35 days after 1^st^ immunization after the 2^nd^ immunization (T3), and 3.5 months (105 ± 7 days) (T4) and 6 months thereafter (185 ± 6 days) (T5) (**Figure 1**). Both cohorts of mRNA vaccinees (mRNA-1273, BNT162b2) received two doses of the vaccine (approximately 28 and 21 days apart, respectively). Ad26.COV2.S was authorized as a 1-dose vaccine and thus blood donation timepoints were based on the initial immunization date. For NVX-CoV2373, volunteers were recruited locally who had participated in a NVX-CoV2373 efficacy trial of two intramuscular 5 μg doses of NVX-CoV2373 plus adjuvant 21 days apart (Dunkle *et al*., 2021). The NVX-CoV2373 trial was structured such that donors initially received two doses of placebo or vaccine in a blinded manner and were then provided two doses of the opposite (vaccine or placebo), such that all participants were vaccinated (Clinicaltrials.gov). Characteristics of the donor cohorts are shown in **Figure 1A-B**. All four vaccine groups were similar in their distribution of gender, age, and race or ethnicity. To measure possible exposure to natural SARS-CoV-2 infection, IgG levels against the Nucleocapsid (N) protein were measured in each vaccinee (**Figure S1A.** See Methods for exclusion criteria).

**Figure 1.**
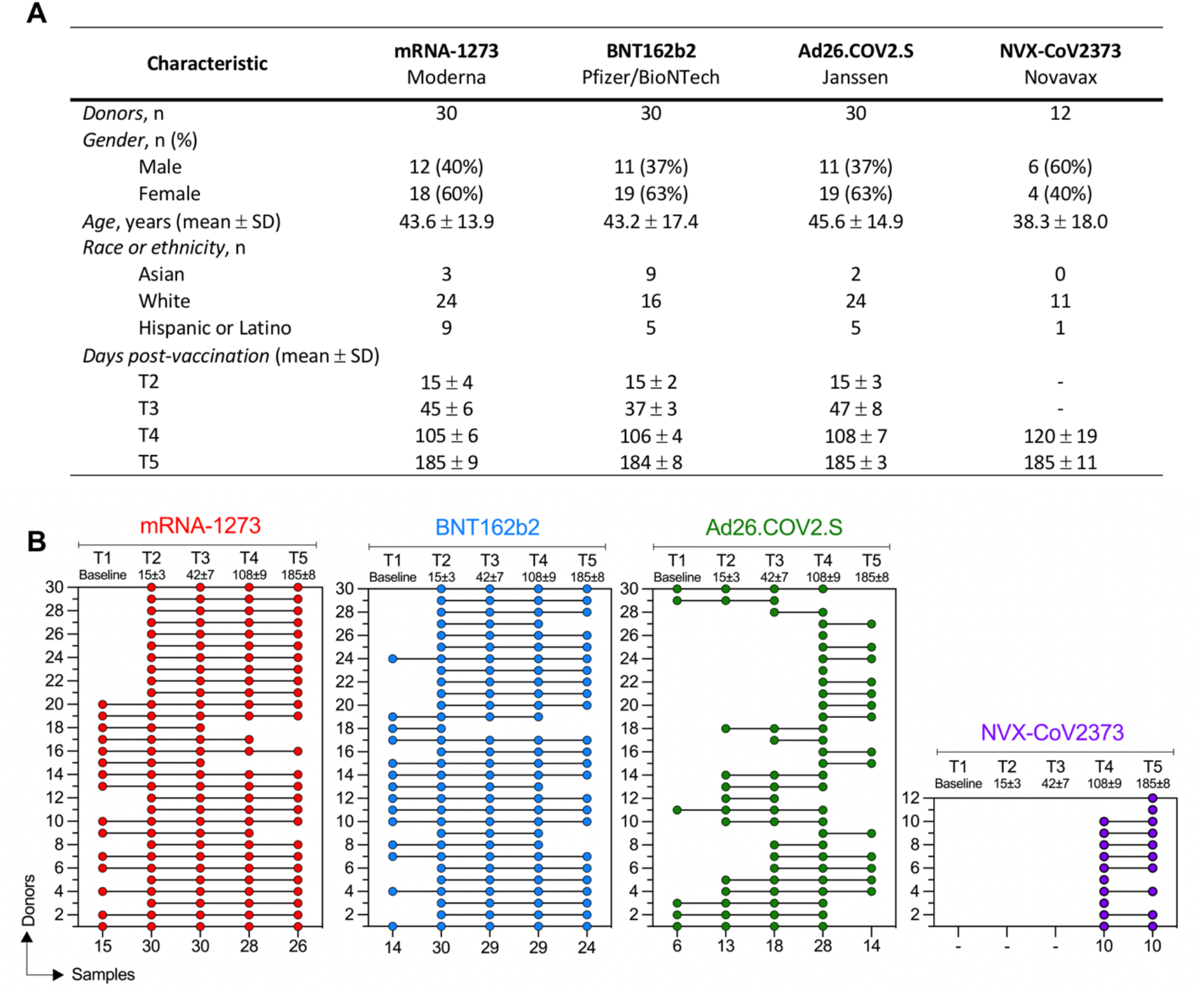
COVID-19 vaccine recipient cohorts. **(A)** Donor cohort characteristics. **(B)** Subjects received mRNA-1273, BNT162b2, Ad26.COV2.S, or NVX-CoV2373 vaccine and donated blood at different times post-vaccination. Each COVID-19 vaccine cohort is color-coded: mRNA-1273 (red), BNT162b2 (blue), Ad26.COV2.S (green), or NVX-CoV2373 (purple). The first column displays the number of donors included in each vaccine cohort and the bottom row shows the number of samples collected for each time point.

### Spike antibody magnitude and durability elicited by different vaccine platforms

For all donors at all available time points, SARS-CoV-2 spike antibodies (**Figure 2A**), receptor-binding domain (RBD) antibodies (**Figure 2B**), N antibodies (**Figure S1A**), and SARS-CoV-2 pseudovirus (PSV) neutralization titers (**Figure 2C**) were determined, for a total of 1,408 measurements from 352 samples. Binding antibody titers and PSV neutralization titers were quantified based on a WHO standard.

**Figure 2.**
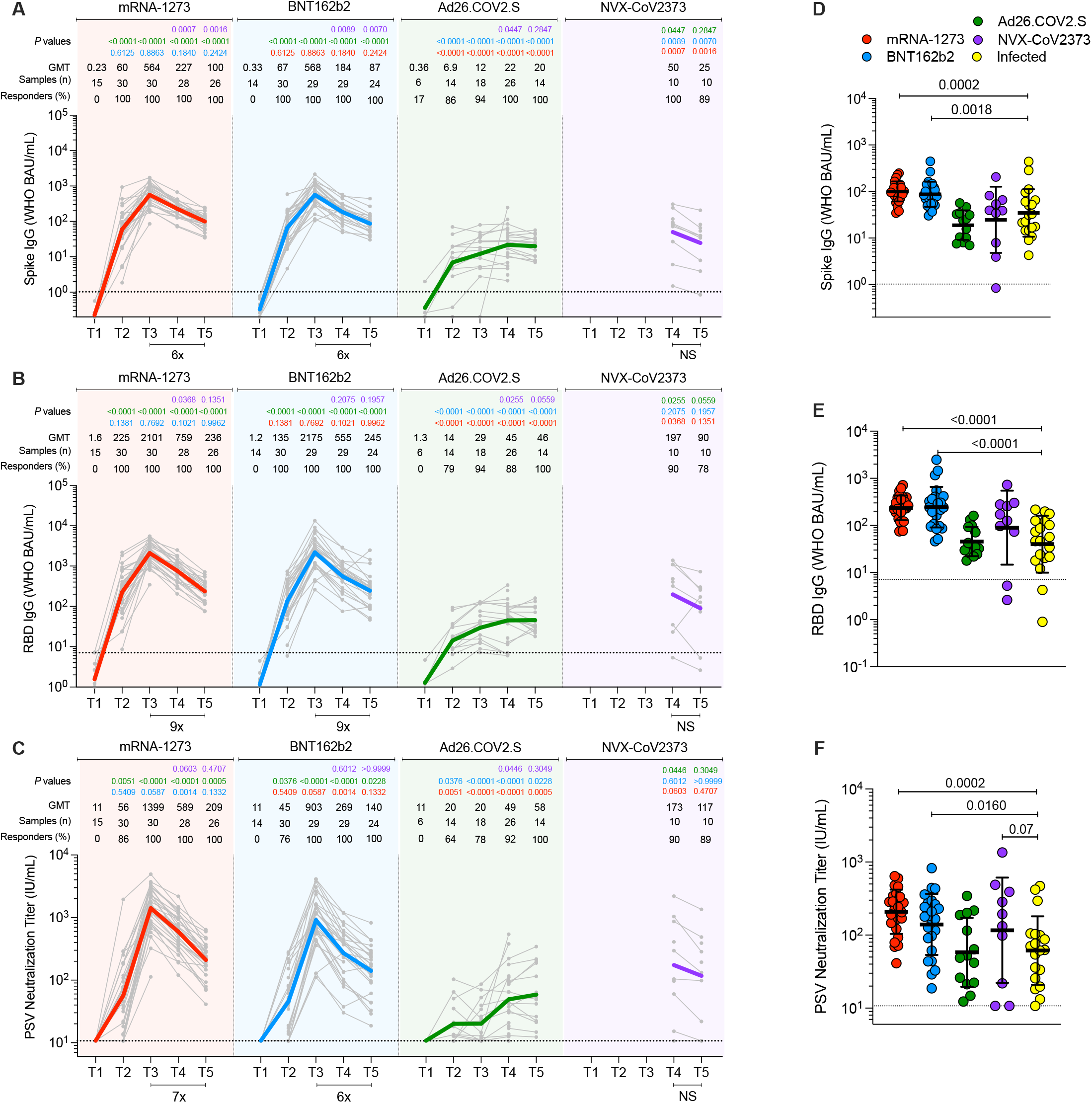
Antibodies elicited by mRNA-1273, BNT162b2, Ad26.COV2.S, and NVX-CoV2373 COVID-19 vaccine platforms. **(A-C)** (A) Comparison of longitudinal SARS-CoV-2 spike IgG levels, (B) SARS-CoV-2 RBD IgG levels, and (C) SARS-CoV-2 pseudovirus neutralizing titers (PSV) from all donors to the mRNA-1273 (red), BNT162b2 (blue), Ad26.COV2.S (green) and NVX-CoV2373 (purple) COVID-19 vaccines over 6 months. Individual subjects are show as gray symbols with connecting lines for longitudinal samples. Geometric means are shown in thick colored lines. Dotted lines indicate the limit of quantification (LOQ). P values show differences between each time point between the different vaccines, color-coded per comparison based on the vaccine compared. NS, nonsignificant; GMT, geometric mean titers. Bottom bars indicate fold changes between two time points. **(D-F)** (D) Comparison of spike IgG, (E) RBD IgG, and (F) PSV neutralization titers at 185 ± 6 days post-vaccination to SARS-CoV-2-infected individuals at 170 to 195 days post-symptom onset. Statistical analysis by Mann-Whitney t-test.

For mRNA-1273, after 1^st^ dose immunization, 100% of vaccinees had detectable spike IgG and RBD IgG titers (**Figures 2A-B**). 86% of vaccinees had detectable neutralization antibody titers after the 1^st^ dose (**Figure 2C**). These early findings are consistent with a large mRNA-1273 clinical trial cohort that measured serology at early time points (100% positive for RBD IgG and spike IgG, 82% positive for neutralization antibody (Gilbert *et al*., 2022). After the 2^nd^ immunization, antibody levels both spike and RBD IgG were boosted 9-fold (**Figures 2A-B**) and neutralization antibody titers were boosted 25-fold (GMT 1,399) (**Figure 2C**). 100% of mRNA-1273 recipients remained positive for spike IgG, RBD IgG, and neutralization antibodies at 6-months post-vaccination (T5). From peak (T3) to 6-months (T5), GMTs of spike IgG decreased 6-fold, RBD IgG decreased 9-fold, and neutralizing antibodies decreased 7-fold.

For BNT162b2, after 1^st^ dose immunization, 100% of vaccinees had detectable Spike IgG and RBD IgG titers (**Figures 2A-B**). 76% of vaccinees had detectable neutralization antibodies after the 1^st^ dose, which was slightly lower than the 86% with mRNA-1273 (**Figure 2C**). After the 2^nd^ immunization, spike and RBD IgG were boosted 9- to 16-fold (**Figures 2A-B**), and neutralization antibodies titers were boosted 20-fold (GMT 903) (**Figure 2C**). 100% of BNT162b2 recipients remained positive for spike IgG, RBD IgG, and neutralization antibodies at 6-month post-immunization (**Figures 2A-C**). From peak (T3) to 6-month (T5), GMT of spike IgG, RBD IgG, and neutralization antibody titers decreased by 6-fold, 9-fold, and 6-fold, respectively. These antibody declines after BNT162b2 immunization were comparable with declines after mRNA-1273 immunization (**Figures 2A-C**). Neutralization antibody titers in BNT162b2 recipients were lower than mRNA-1273 recipients by 1.6-fold (p=0.059), 2.2-fold (p=0.0014), and 1.5-fold (p=0.13), at the T3, T4, and T5 time points, respectively. Neutralization antibody titers trended lower in BNT162b2 than mRNA-1273 recipients when assessed in aggregate across the entire 6-month time period (area under curve (AUC), p=0.051, **Figures S1B-D**).

For Ad26.COV2.S 1-dose immunization, 86% of vaccinees had detectable Spike IgG and 79% RBD IgG at T2 (**Figures 2A-B**). 64% of vaccinees had detectable neutralization antibodies at T2, which was somewhat lower than the 86% with mRNA-1273 and 76% with BNT162b2 (**Figure 2C**). Ad26.COV2.S antibody binding and neutralization titers gradually increased over time, with 100% of recipients having detectable Spike IgG, RBD IgG, and neutralization antibodies at 6-month post-immunization. Ad26.COV2.S neutralization antibody titers peaked at T5 (GMT 58), but that peak was still 24-fold lower than the mRNA-1273 peak (GMT 1,399) and 16-fold lower than the BNT162b2 peak (GMT 903). At 6-month post-immunization, Ad26.COV2.S neutralization antibody titers were 3.6-fold lower than mRNA-1273 and 2.4-fold lower than BNT162b2 (**Figure 2C**). Over the entire 6-month time period, Ad26.COV2.S spike IgG, RBD IgG, and neutralization antibody titers were significantly lower than mRNA vaccine recipients (p<0.0001 mRNA-1273, p<0.0001 BNT162b2. **Figures S1B-D**).

For NVX-CoV2373, antibody titers were available for 3.5 and 6 months. Spike and RBD IgG titers were substantial at 3.5 months post-vaccination and were marginally (not significantly) decreased at T5 (**Figures 2A-B**). Neutralization antibody titers were comparable at both timepoints (**Figure 2C**). At 6-month post-immunization, NVX-CoV2373 neutralization antibody titers (GMT 152) were 2.6 fold higher than Ad26.COV2.S (GMT 58), and were comparable to mRNA-1273 (GMT 209) and BNT162b2 (GMT 140). Considering the 3.5-month to 6-month period in aggregate, RBD IgG and neutralization antibody titers in NVX-CoV2373 recipients were comparable to both mRNA vaccines (**Figures S1F-G**).

Lastly, antibody titers at 6 months were compared to SARS-CoV-2 infected subjects (**Figures 2D-F**) who were enrolled for a previously reported study (Mateus et al., 2021). The previously infected individuals were selected randomly. Recipients of the mRNA vaccines (mRNA-1273 and BNT162b2) had 4.5-fold higher spike IgG (**Figure 2D**), 6.4-fold higher RBD IgG (**Figure 2E**), and 3.4-fold higher neutralization antibody titers (**Figure 2F**) compared to previously-infected subjects. Antibody titers from NVX-CoV2373 recipients also trended higher than SARS-CoV-2 infected subjects (**Figures 2D-F**). Antibody titers from Ad26.COV2.S were similar to titers from SARS-CoV-2 infected subjects (**Figures 2D-F**).

Overall, antibody titers were significantly higher for mRNA recipients than Ad26.COV2.S recipients. Recipients of NVX-Co2373 immunization also had higher peak antibody titers than recipients of Ad26.COV2.S. Antibody titers to mRNA-1273, BNT162b2, and Ad26.COV2.S changed substantially over the 6+ months of observation, with different patterns seen for the mRNA versus adenoviral vector platforms.

### Spike-specific CD4^+^ T cell memory elicited by four different vaccines

SARS-CoV-2 spike-specific CD4^+^ T cell responses were measured for all donors at all available timepoints utilizing two previously described flow cytometry activation-induced marker (AIM) assays (OX40^+^CD137^+^ and OX40^+^ surface CD40L^+^ (sCD40L)) (**Figures 3A, 3D,** and **S2-S3**) and separate intracellular staining (ICS) for cytokines (IFNγ, TNFα, IL-2), granzyme B (GzB), and intracellular CD40L (iCD40L) (**Figures 3C, 3F, 4** and **S4**). SARS-CoV-2 spike-specific circulating follicular helper T (cTfh) cells were measured at all time points (**Figures 3B, 3E, S2A** and **S2D**), as this subpopulation of CD4^+^ T cells is crucial for supporting antibody responses following vaccination (Crotty, 2019; Lederer et al., 2022; Mudd et al., 2022).

**Figure 3.**
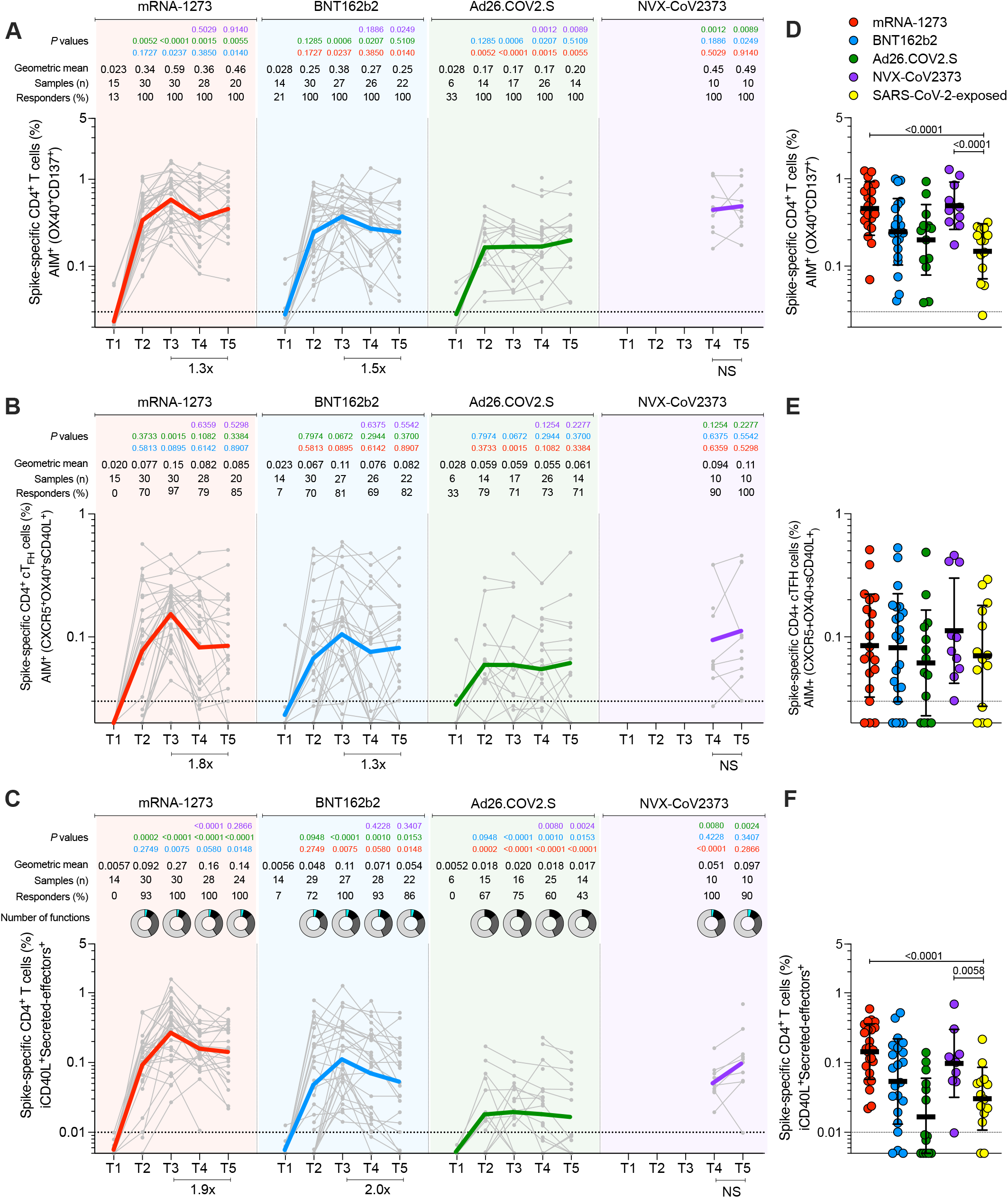
Acute and memory CD4^+^ T cell responses after mRNA-1273, BNT162b2, Ad26.COV2.S, or NVX-CoV2373 immunization. **(A)** Longitudinal spike-specific CD4^+^ T cell responses induced by four different COVID-19 vaccines measured by OX40^+^ CD137^+^ AIM after spike megapool (MP) stimulation. See **Figures S2B-C** for the representative gating strategy of AIM^+^ cells. **(B)** Longitudinal spike-specific circulating T follicular helper cells (cTfh) induced by COVID-19 vaccines. Spike-specific cTfh cells (CXCR5^+^OX40^+^sCD40L^+^, as % of CD4^+^ T cells) after stimulation with spike MP. See **Figure S2D** for representative gating strategy. **(C)** Spike-specific CD4^+^ T cells measured by ICS. Expressing iCD40L and producing IFNγ, TNFα, IL-2, or GzB (Secreted-effector^+^ = ICS^+^). See **Figure 4** for analysis of individual IFNγ, TNFα, IL-2, or GzB on spike-specific CD4^+^ cytokine^+^ T cells expressing iCD40L). Donut charts depict the proportions of multifunctional secreted effector profiles among the spike-specific ICS^+^ CD4^+^ T cells: 1 (light gray), 2 (dark gray), 3 (black), and 4 (turquoise) functions (See also **Figure S4**). **(D-F)** Comparison of spike-specific CD4^+^ T cells by AIM (D), cT_FH_ (E), and ICS (F) between COVID-19 vaccinees at 185 ± 6 days post-vaccination and SARS-CoV-2-exposed subjects 170 to 195 days PSO. The dotted black line indicates the limit of quantification (LOQ). The color-coded bold lines in (A), (C), and (E) represent the geometric mean in each time post-vaccination. Background-subtracted and log data analyzed. P values on the top in (A), (C), and (E) show the differences between each time point in the different vaccines and are color-coded as per Figure 1. Bottom bars in (A), (C), and (E) show fold-changes between T3 and T5. Data were analyzed for statistical significance using the Mann-Whitney test [(A-F)]. NS, nonsignificant.

**Figure 4.**
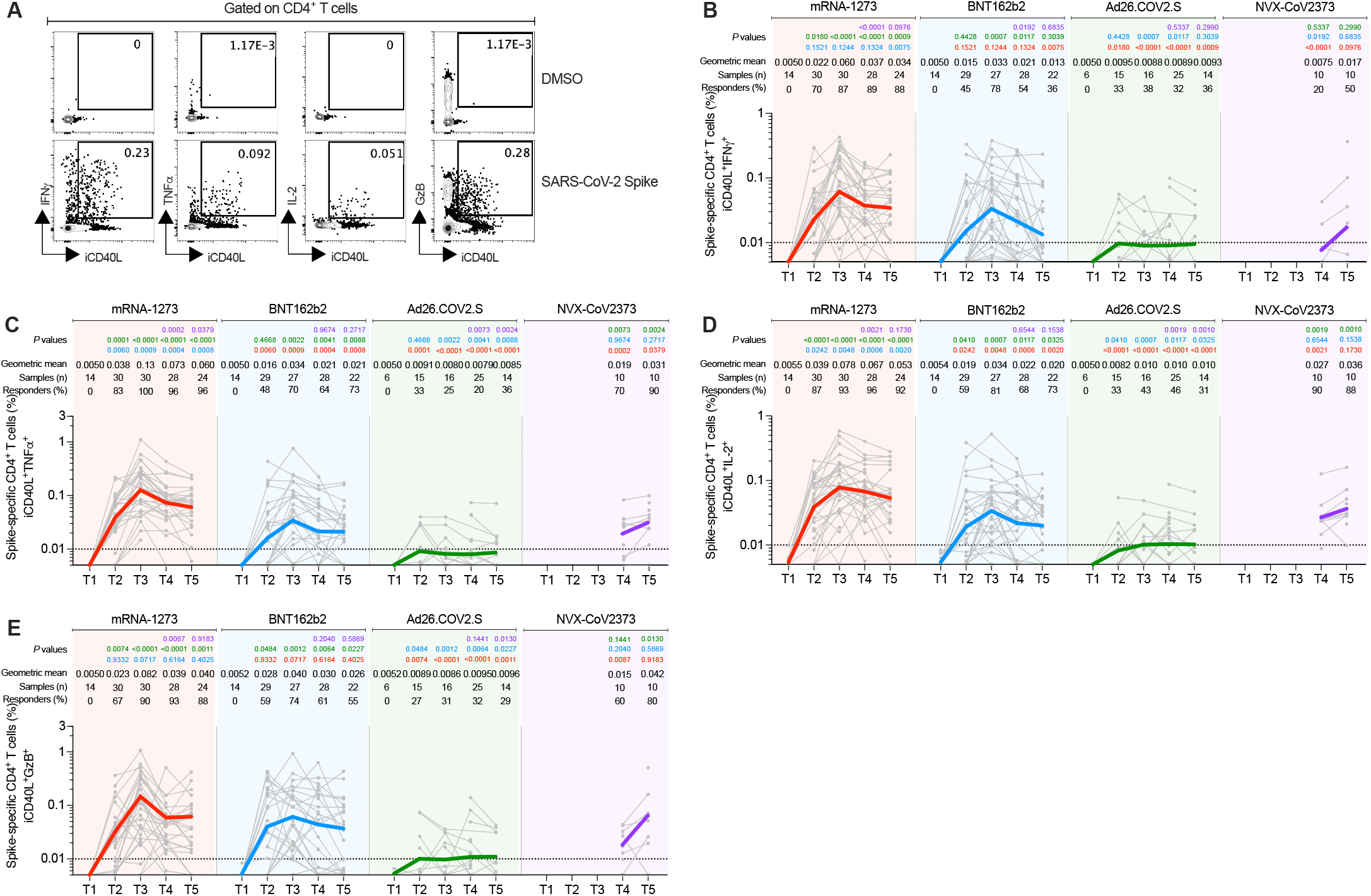
Longitudinal spike-specific CD4^+^ T cells expressing intracellular CD40L (iCD40L) and producing cytokines or granzyme B in subjects vaccinated with the mRNA-1273, BNT162b2, Ad26.COV2.S, or NVX-CoV2373 COVID-19 vaccines. **(A)** Representative gating strategy of spike-specific CD4^+^ T cells expressing iCD40L^+^ producing cytokines or Granzyme B (GzB) detected in COVID-19 vaccine platforms at T3. Secreted-effector^+^ CD4^+^ T cell responses were quantified by expressing iCD40L^+^ along with the production of IFNγ, TNFα, IL-2, and/or GzB after stimulation with spike megapool (MP). **(B-E)** Spike-specific CD4^+^ T cells expressing iCD40L^+^ and producing IFNγ(B), TNFα (C), IL-2 (D), or GzB (E) from COVID-19 vaccinees evaluated at T1, T2, T3, T4, and T5. The dotted black line indicates the limit of quantification (LOQ). The color-coded bold lines in (B-E) represent the Geometric mean in each time post-vaccination. Background-subtracted and log data analyzed. P values on the top in (B-E) show the differences between each time point in the different vaccines and are color-coded as follows: mRNA-1273 (red), BNT162b2 (blue), Ad26.COV2.S (green), or NVX-CoV2373 (purple). Data were analyzed for statistical significance using the Mann-Whitney test [(B-E)]. T1, Baseline; T2, 15 ± 3 days; T3, 42 ± 7 days; T4, 108 ± 9 days; T5, 185 ± 8 days.

In response to a single dose of the mRNA-1273 vaccine (T2), a majority of subjects developed a spike-specific CD4^+^ T cell response as measured by both AIM^+^ (**Figures 3A** and **S3**) and iCD40L^+^secreted-effector^+^ (ICS^+^) CD4^+^ T cells (**Figure 3C**). Spike-specific CD4^+^ T cell responses peaked after the 2^nd^ mRNA-1273 vaccination (100% responders, T3) and were well maintained out to 6 months post-vaccination, with only a 1.0- to 1.9-fold reduction in AIM^+^ or ICS^+^ CD4^+^ T cells, respectively (**Figures 3A,C** and **S3**). mRNA-1273 vaccination induced spike-specific cTfh cells in most donors after the 1^st^ dose, which peaked after the 2^nd^ dose (97%, T3), and memory cTfh cells were maintained out to 6 months post-vaccination with only a 1.4-fold change from peak (T3 to T5, **Figure 3B**). Memory cTfh cells represented 27% of the spike-specific memory CD4^+^ T cells, on average.

Vaccination with BNT162b2 induced spike-specific AIM^+^ and ICS^+^ CD4^+^ T cells after the first vaccination (T2), with peak responses after the 2^nd^ immunization (T3) (**Figures 3A, 3C** and **S3**). However, peak responses to BNT162b2 vaccination were significantly lower than mRNA-1273 peak vaccine responses both by AIM and ICS (1.6-fold lower, P=0.019; and 2.5-fold lower, P=0.011. **Figures 3A** and **C**). Memory CD4^+^ T cells were detectable in 85-100% of BNT162b2 vaccinees at 6 months after immunization, but the memory CD4^+^ T cell frequencies were significantly lower than for mRNA-1273 (1.9-fold lower by AIM, P=0.011 and 2.4-fold lower by ICS, P=0.038, **Figures 3A** and **3C**). Spike-specific memory cTfh cell frequencies were comparable between BNT162b2 and mRNA-1273 vaccination (**Figure 3B**).

Both mRNA-1273 and BNT162b2 vaccination induced ICS^+^ spike-specific memory CD4^+^ T cells, including iCD40L^+^IFNγ^+^, iCD40L^+^ TNFα^+^, and iCD40L^+^IL-2^+^ cells, detectable out to 6 months post-vaccination. mRNA-1273 vaccinees had significantly higher frequencies of TNFα^+^ and IL-2^+^ CD4^+^ T cells at all timepoints and higher levels of IFNγ^+^ memory CD4^+^ T cells at 6 months relative to BNT162b2 vaccinees (**Figure 4**). GzB^+^ CD4^+^ T cells (iCD40L^+^GzB^+^) were assessed as indicators of CD4^+^ cytotoxic T lymphocytes (CD4-CTL). Interestingly, both mRNA vaccines generated CD4-CTLs as a significant fraction of the overall spike-specific CD4^+^ T cell response (**Figures 4E** and **S4B**). Multifunctional spike-specific CD4^+^ T cells were observed after the 1^st^ dose of either mRNA-1273 or BNT162b2, and multifunctionality was stably maintained out to 6 months (**Figures 3C** and **S4A**).

For the Ad26.COV2.S vaccine, spike-specific CD4^+^ T cell responses were detectable in a majority of individuals and were largely stable out to 6 months post-vaccination (69-100% of individuals with spike-specific CD4^+^ T cells by AIM assays; 46% with spike-specific CD4^+^ T cells by ICS. **Figures 3A, 3C**, and **S3**). cTfh cells were detectable in the majority of individuals (**Figure 3B**). Peak CD4^+^ T cell responses were lower to Ad26.COV2.S than either of the mRNA vaccines. Peak AIM^+^ CD4^+^ T cells to Ad26.COV2.S were 2.2- to 3.3-fold lower than BNT162b2 and 3.5- to 4.2-fold lower than mRNA-1273 peak responses (**Figures 3A** and **S3**). Peak spike-specific ICS^+^ CD4^+^ T cell responses to Ad26.COV2.S were 5.8-fold lower than BNT162b2 and 14-fold lower than mRNA-1273 (**Figure 3C**). Both mRNA vaccines generated significantly higher peak frequencies of IFNγ^+^ CD4^+^ T cells than Ad26.COV2.S vaccination (iCD40L^+^IFNγ^+^, mRNA1273 P<0.0001, BNT162b2 P=0.001), and mRNA-1273 vaccinees had significantly higher IFNγ^+^ spike-specific memory CD4^+^ T cells than Ad26.COV2.S at 6 months post-vaccination (P=0.007, **Figure 4**). The mRNA vaccines also induced significantly more CD4-CTLs at peak than Ad26.COV2.S (mRNA1273 P<0.0001, BNT162b2 P=0.0012, **Figure 4E**), and the CD4-CTLs induced by the mRNA vaccines were more sustained as memory cells at the 6-month memory timepoint relative to Ad26.COV2.S (**Figure 4E**). Spike-specific CD4^+^ T cells induced by Ad26.COV2.S had less multifunctionality at all time points relative to both mRNA vaccines (**Figures 3E** and **S4A**). Overall, memory CD4^+^ T cell frequencies were lower after Ad26.COV2.S immunization compared to mRNA vaccines, assessed as total spike-specific memory (AIM^+^), cTfh memory, IFNγ^+^ memory, CD4-CTL memory, or memory CD4^+^ T cell multifunctionality.

For the NVX-CoV2373 vaccine, 100% of immunized individuals developed spike-specific memory CD4^+^ T cells detected by both AIM and ICS assays (**Figures 3A, 3C** and **S3**). All NVX-CoV2373 immunized individuals had spike-specific memory cTfh cells (**Figure 3B**). Memory CD4^+^ T cell responses to NVX-CoV2373 were comparable in magnitude to the mRNA vaccines by AIM (**Figures 3A, 3C** and **S3**). By ICS, NVX-CoV2373 responses 6 months post-vaccination were comparable to BNT162b2 (NVX-CoV2373 geomean 0.074%, BNT162b2 0.059%), and significantly higher than the Ad26.COV2.S vaccine (Ad26.COV2.S geomean 0.015%, P=0.0057. **Figure 3C**). NVX-CoV2373 induced multifunctional memory spike-specific CD4^+^ T cells comparably to both mRNA vaccines (T4 and T5, **Figures 3C** and **S4A**), with a shift in the relative abundance of IL-2^+^ cells over IFNγ^+^ memory CD4^+^ T cells observed for NVX-CoV2373 (**Figures 4B** and **4D**).

Spike-specific CD4^+^ T cell responses in COVID-19 recovered individuals were assessed to compare infection-induced versus vaccine-elicited T cell memory (**Figures 3D-F**). Spike-specific CD4^+^ T cell memory at 6 months post-vaccination in mRNA-1273 and NVX-CoV2373 vaccinees was significantly higher than for COVID-19 recovered individuals, both by AIM and ICS (**Figures 3D** and **3F**). BNT162b2 and Ad26.COV2.S generated memory CD4^+^ T cells frequencies not significantly different than SARS-CoV-2 infection (**Figures 3D** and **3F**). Memory cTfh cell frequencies were similar between all four vaccines and infection (**Figure 3E**). Overall, all four of the COVID-19 vaccines generated memory CD4^+^ T cells in the majority of vaccinated individuals, with representation of both Th1 (IFNγ^+^) and Tfh memory, with memory CD4-CTL also generated by mRNA and NVX-CoV2373 vaccines. Additionally, the magnitude of spike-specific CD4^+^ T cell memory was generally higher for mRNA vaccines and NVX-CoV2373 than seen in COVID-19 recovered individuals.

### Spike-specific CD8^+^ T cells elicited by four different vaccines

SARS-CoV-2 spike-specific CD8^+^ T cells were measured by ICS at all time points to identify IFNγ, TNFα, or IL-2 producing cells (CD69^+^ cytokine^+^ gating = “ICS^+^”. **Figures 5A-C, S5-7**) for all vaccine modalities. Spike-specific CD8^+^ T cells were also measured by AIM (CD69^+^CD137^+^, **Figure S8**).

**Figure 5.**
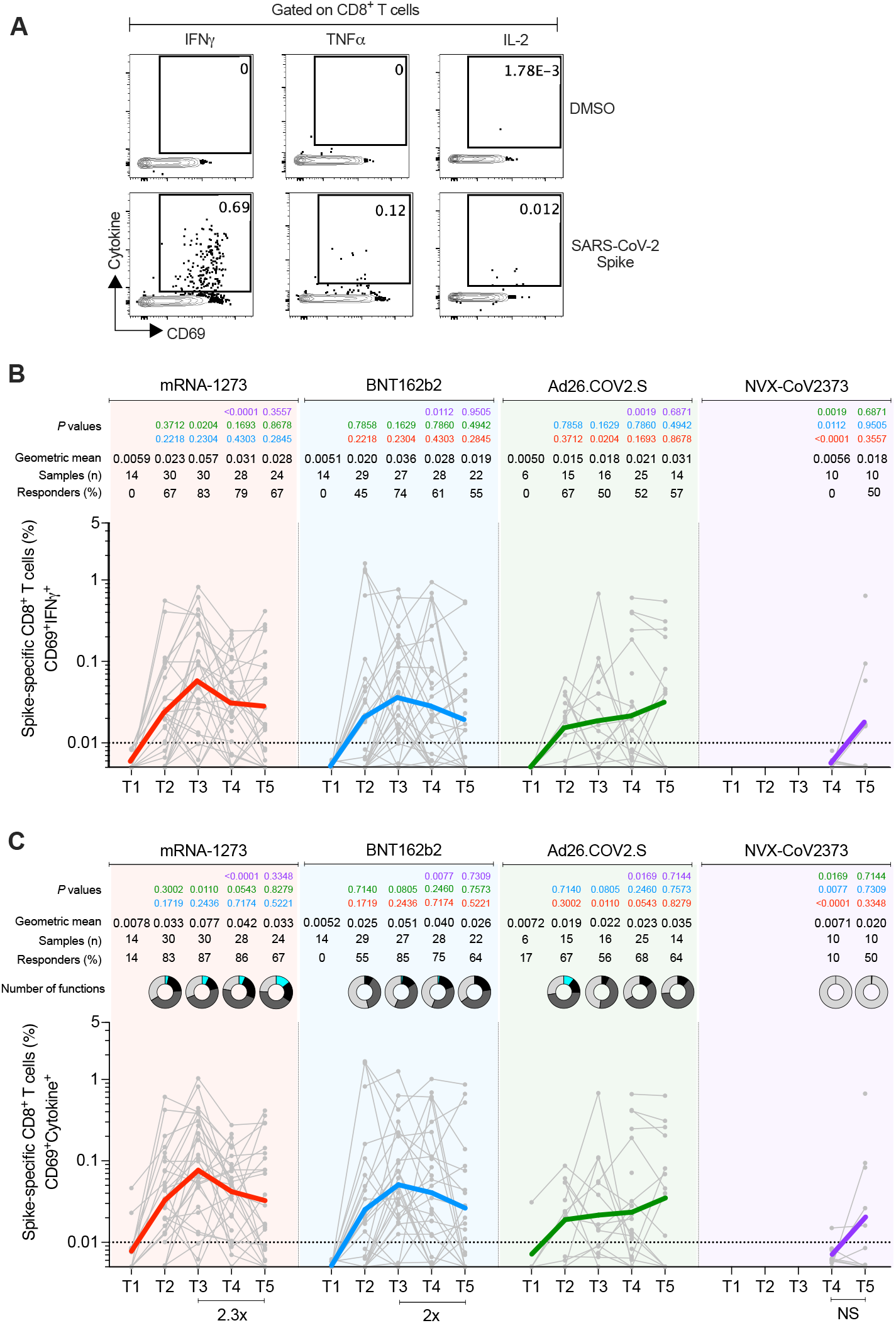
Acute and memory CD8^+^ T cell responses after mRNA-1273, BNT162b2, Ad26.COV2.S, or NVX-CoV2373 immunization. **(A)** Representative gating of spike-specific CD8^+^ T cells. Cytokine-producing (“cytokine^+^”) CD8^+^ T cells were quantified as CD69^+^ along with IFNγ, TNFα, or IL-2 expression after stimulation with spike MP. **(B)** Longitudinal quantitation of CD69^+^IFNγ^+^ spike-specific CD8^+^ T cells. See **Figure S5** for TNFα and IL-2, and **Figure S9** for additional analysis. **(C)** Longitudinal quantitation of cytokine^+^ spike-specific CD8^+^ T cells. CD8^+^ T cells were quantified as CD69^+^ along with IFNγ, TNFα, or IL-2 expression after stimulation with spike MP. Bottom bars show fold-changes between T3 and T5. The donut charts depict the proportions of multifunctional cytokine^+^ profiles of the spike-specific CD8^+^ T cells, including IFNγ, TNFα, or IL-2 and GzB: 1 (light gray), 2 (dark gray), 3 (black), and 4 (turquoise) functions (See also **Figure S9**). The dotted black line indicates the limit of quantification (LOQ). Graphs are color-coded as per Figure 2. Background-subtracted and log data analyzed. Data were analyzed for statistical significance using the Mann-Whitney test [(B), (C)].

For the mRNA-1273 vaccine, 83% of vaccinees had detectable spike-specific CD8^+^ T cell responses after the 1^st^ immunization (**Figure 5C**). ICS^+^ CD8^+^ T cell response rates peaked after the 2^nd^ immunization (87% T3 responders **Figure 5C**). Spike-specific memory CD8^+^ T cells were largely maintained out to 6 months after mRNA-1273 vaccination (67% responders, **Figure 5C**), with only a 2.3-fold decline in geomean frequency from the peak (0.077% to 0.033%, **Figure 5C**). Both acute and memory CD8^+^ T cell responses were dominated by IFNγ-producing cells (**Figures 5B-C** and **S9**), the majority of which co-expressed GzB (**Figure S9**). The majority of the memory spike-specific CD8^+^ T cells exhibited an effector memory (T_EM_) surface phenotype (**Figure S10**).

For the BNT162b2 vaccine, IFNγ^+^ and total ICS^+^ CD8^+^ T cell responses also peaked after the 2^nd^ immunization (T3 73% and 85% responders, respectively **Figures 5B-C**). Memory CD8^+^ T cells were maintained out to 6 months after BNT162b2 vaccination (60% responders, **Figure 5C**), with only a 1.8-fold decline in geomean frequency (**Figure 5C**). Multifunctional spike-specific memory CD8^+^ T cells were more common in mRNA-1273 compared to BNT162b2 vaccinees (**Figures 5C** and **S9A**), with the responses dominated by IFNγ^+^ cells (**Figures 5C,** and **S9**). Overall, spike-specific CD8^+^ T cell acute and memory responses to BNT162b2 were similar to mRNA-1273 but slightly lower in frequency and multifunctionality.

The fraction of CD8^+^ T cell responders to Ad26.COV2.S was lower than both mRNA vaccines (71% compared to 87% and 85%, **Figure 5C**). Nevertheless, Ad26.COV2.S spike-specific CD8^+^ T cell frequencies were relatively stable through 6 months post-vaccination (**Figures 5B-C** and **S5**) and geomean frequencies of memory CD8^+^ T cells after Ad26.COV2.S vaccination were comparable to both mRNA vaccines at 6 months (**Figures 5B-C** and **S5**).

For the NVX-CoV2373 vaccine, spike-specific ICS^+^ memory CD8^+^ T cells were observed in 10% to 40% of donors (T4 and T5, **Figure 5C**). There were minimal multifunctional CD8^+^ T cells (**Figure 5C** and **S9**).

Overall, memory CD8^+^ T cell frequencies and response rates were similar between mRNA-1273, BNT162b2, and Ad26.COV2.S immunizations. Low but detectable memory CD8^+^ T cells were observed in some individuals after NVX-CoV2373 immunization. CD8^+^ T cell responses to all COVID-19 vaccines were dominated by IFNγ-producing cells. No differences in IFNγ MFI were observed between memory CD8^+^ T cells generated to each of the vaccines (**Figure S7**). All vaccines elicited AIM^+^ (CD69^+^CD137^+^) CD8^+^ cell responses at levels comparable to, or slightly higher than, frequencies observed in SARS-CoV-2 recovered individuals. (**Figure S6**).

### Spike- and RBD-specific B cell memory to four COVID-19 vaccines

Next, we sought to characterize and compare the development of B cell memory across the 4 different COVID-19 vaccines. For that, we utilized spike and RDB probes to identify, quantify and phenotypically characterize memory B cells from vaccinated subjects at 3.5 (T4) and 6 months (T5) after immunization (**Figures 6A-B** and **S11**). Spike-specific and RBD-specific memory B cells were detected in all vaccinated subjects at 6 months (**Figures 6C-D**). RBD-specific memory B cells comprised 15 to 20% of the spike-specific memory B cell population, on average (**Figure S12A**). Immunization with mRNA-1273 or BNT162b2 led to higher frequencies of spike-specific and RBD-specific memory B cells compared to Ad26.COV2.S and NVX-CoV2373 at 3.5 and 6 months (each p<0.01. **Figures 6C-D**).

**Figure 6.**
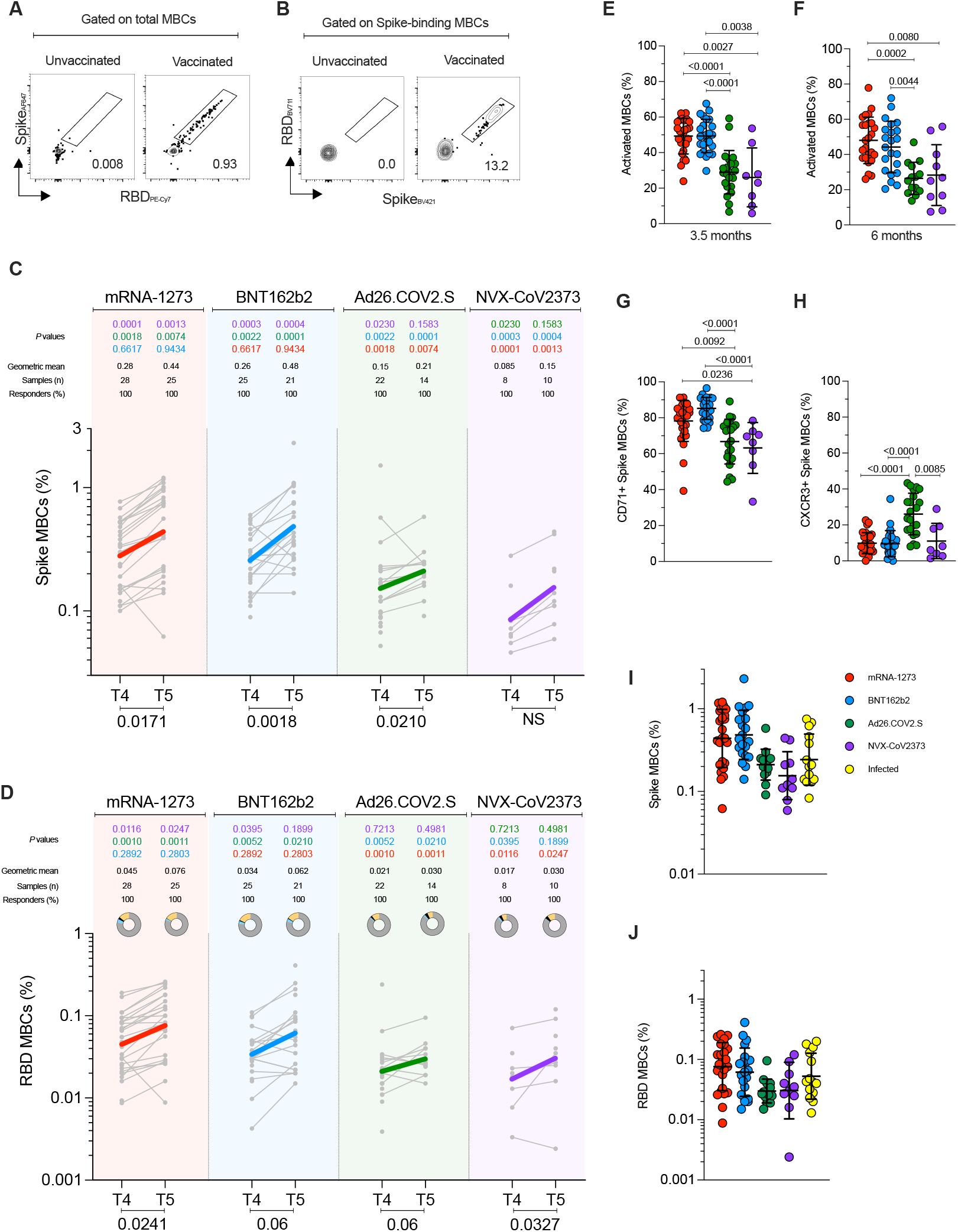
SARS-CoV-2-specific memory B cells to mRNA-1273, BNT162b2, Ad26.COV2.S, and NVX-CoV2373 vaccines. **(A-B)** Representative gating strategy for **(A)** spike-binding and **(B)** RBD-binding memory B cells (“MBCs”) (See also **Figure S11**). **(C-D)** Frequency of **(C)** spike-binding and **(D)** RBD-binding MBCs from total MBCs elicited after 3.5 and 6 months. Limit of detection= 0.0017. RBD donut graphs represent isotype distribution; IgG (grey), IgA (blue), IgM (black), and other (yellow). **(E and F)** Proportion of spike-binding MBCs with activated phenotype (CD21-CD27+) at **(E)**3.5 and **(F)**6 months. **(G and H)** Proportion of spike-binding MBCs expressing **(G)** CD71 or **(H)** CXCR3 at 3.5 months. **(I and J)** Comparisons between vaccinees and SARS-CoV-2-infected individuals for (I) Spike-binding MBCs and (J) RBD-binding MBCs at 6 months. The vaccines are color-coded as per Figure 2. The color-coded bold lines in (B) and (D) represent the geometric mean at each time post-vaccination. Bottom bars show T4 to T5 statistics. Data were analyzed for statistical significance using the Mann-Whitney test [(B), (D)], Kruskal-Wallis (KW) test and Dunn’s post-test for multiple comparisons [(E), (F), (G), (H), (I), (J)]. NS, non-significant. See also **Figure S12**.

Memory B cell responses to the 4 vaccines did not exhibit the same kinetics as the antibody responses. The frequency of spike-specific memory B cells increased over time, (mRNA-1273, p=0.017; BNT162b2, p=0.0018, Ad26.COV2.S, p=0.021. **Figure 6C**). RBD-specific memory B cell frequencies increased at 6 months after mRNA-1273 (1.7-fold, p=0.024), BNT162b2 (2.2-fold, p=0.06), Ad26.COV2.S (2.1-fold, p=0.06), and NVX-CoV2373 (3.05-fold, p=0.033) (**Figure 6D**).

RBD-specific memory B cell isotypes were mostly comparable among the different vaccines, with an average distribution of 83.0% IgG, 2.5% IgM, and 2.2% IgA at 6 months (**Figure 6D** and **Figure S12B**); however, IgA^+^ RBD-specific memory B cells were significantly higher at 3.5 months in mRNA vaccinees compared to Ad26.COV2.S (mRNA-1273 p=0.003. BNT162b2 p=0.04. **Figure 6D**). Phenotypically, activated memory B cells (CD21^-^CD27^+^) comprised 77-85% of spike-specific memory B cells after mRNA vaccination (**Figure 6E** and **S12C**), which was significantly higher than observed for Ad26.COV2.S or NVX-CoV2373 (66%, 61%, mRNA vs. Ad26.COV2.S, p<0.0001. mRNA-1273 p=0.0027; BNT162b2 p=0.0038. **Figure 6E**), and the differences persisted at 6 months (**Figure 6F**). Reciprocally, the representation of classical memory B cells (CD21^+^CD27^+^) was lower in response to mRNA vaccines (**Figure S12D**). To further qualitatively compare memory B cells across vaccine platforms, we assessed CD71, CXCR3, CD95, and CD11c expression by spike-specific memory B cells. CD71^+^ memory B cells were more common at 3.5 months in response to mRNA vaccines than Ad26.COV2.S or NVX-CoV2373 (T4 **Figure 6G**), with higher expression on activated memory B cells (**Figure S12E**). Considering that CD71 is a proliferation marker of B cells, this may reflect greater continuing production of memory B cells in response to mRNA vaccines at 3.5 months compared to Ad26.COV2.S and NVX-CoV2373 vaccines. At 6 months, the frequency of CD71^+^ spike-specific memory B cells remained elevated for mRNA-1273 (**Figure S12F**). CXCR3^+^ spike-specific memory B cell frequencies were substantially higher in response to Ad26.COV2.S compared to the other vaccine platforms (mRNA-1273 p<0.001, BNT162b2 p<0.001, NVX-CoV2373 p=0.008. **Figure 6H**) and remained elevated at 6 months (**Figure S12G**).

Lastly, the frequencies of spike-specific and RBD-specific memory B cells at 6 months post-vaccination were comparable to the frequencies found in previously-infected subjects at 6 months (**Figures 6I-J**), indicating robust memory B cell development to each of the four COVID-19 vaccines.

### Multiparametric comparisons across vaccine platforms

We performed multiparametric analyses, utilizing both correlation matrixes and principal component analysis (PCA) to assess the relative immunogenicity of the four vaccines. Considering all parameters of vaccine antigen-specific immune responses at 6 months after mRNA (mRNA-1273 and BNT162b2) or Ad26.COV2.S vaccination (**Figures S13A-B**), we observed strong correlations between spike IgG, RBD IgG, and neutralization antibody titers (**Figures 7A-B** and **7F**). Neutralization antibody titers correlated with spike-specific and RBD-specific memory B cells for mRNA vaccinees at 6 months (**Figures 7A, C-D**). Antibody levels and memory CD4^+^ T cells were significantly associated in mRNA vaccinees by multiple metrics (**Figures 7A** and **7E**). In contrast, no relationship was observed between antibodies and memory CD8^+^ T cells (**Figure 7A**). Memory CD4^+^ T cells and CD8^+^ T cells were significantly cross-correlated in mRNA vaccinees (**Figure S13A**). For Ad26.COV2.S vaccination, no significant correlations were detected at 6 months between antibodies, memory B cells, memory CD4^+^ T cells, or memory CD8^+^ T cells, which may be related to the smaller cohort size (**Figures 7A** and **7G-I**).

**Figure 7.**
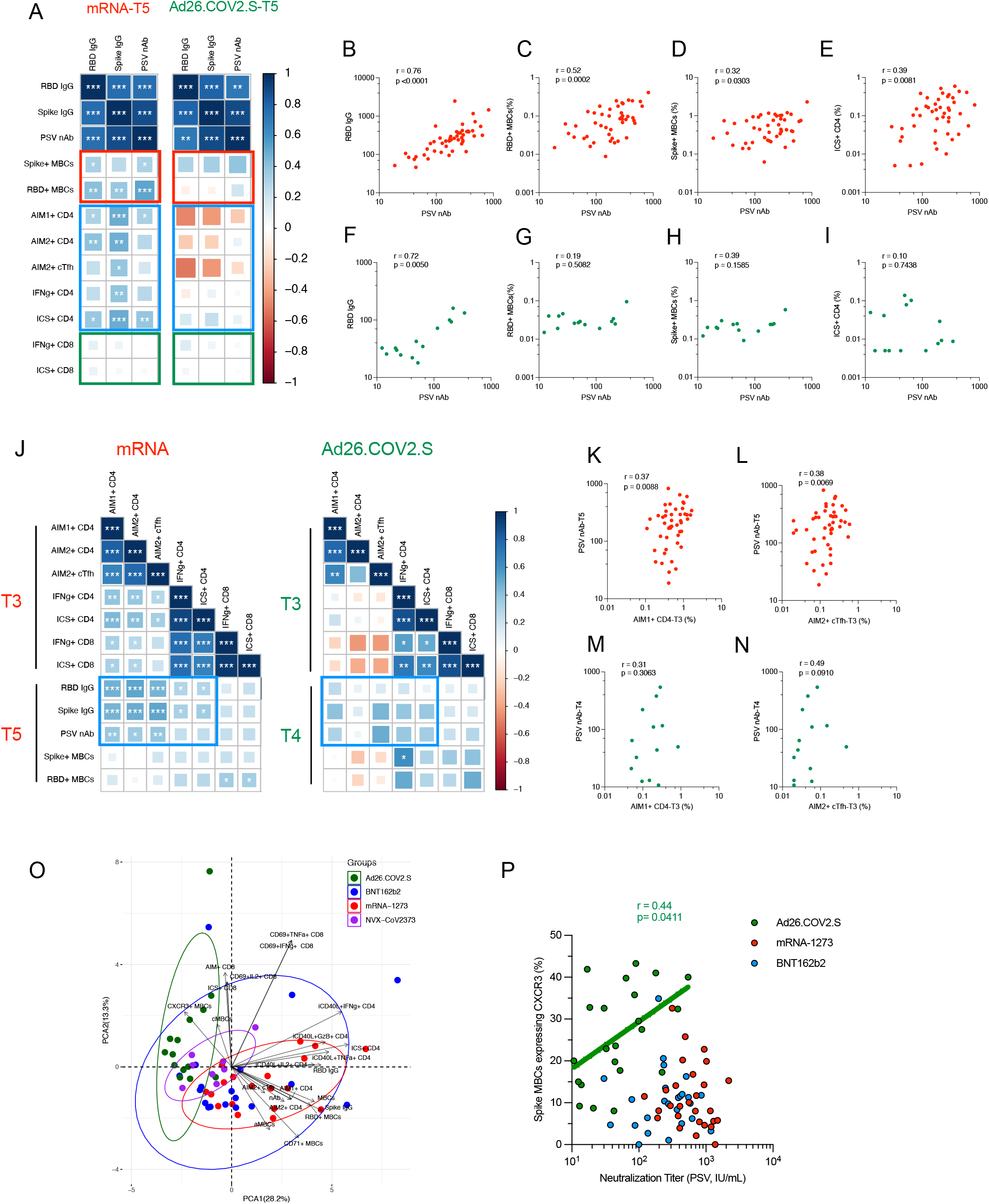
Vaccine-specific correlation analyses. **(A)** Correlation matrix of T5 (6-month) samples, plotted as mRNA (mRNA-1273 and BNT162b2) and Ad26.COV2.S COVID-19 vaccines. The red rectangle indicates the association between antibody and MBC; the blue rectangle indicates the association between antibody and CD4^+^ T cells; the green rectangle indicates the association between antibody and CD8^+^ T cells. Spearman rank-order correlation values (r) are shown from red (−1.0) to blue (1.0); r values are indicated by color and square size. p values are indicated by white asterisks as * p <0.05, **p <0.01, *** p <0.001. MBCs indicates memory B cell, AIM1 indicates OX40^+^CD137^+^, AIM2 indicates OX40^+^sCD40L^+^, nAb indicates neutralization antibody. **(B-I)** The association of indicated parameters shown by scatter plot. Red indicated mRNA, green indicated Ad26.COV2.S. Spearman rank-order correlation values (r) and p values were shown. **(J)** Correlation matrix of CD4^+^ and CD8^+^ T cell data from the early time point with MBCs and antibody data from the late timepoint. The blue rectangle indicates the association between CD4^+^ T cell and antibody. Spearman rank-order correlation values (r) are shown from red (−1.0) to blue (1.0); r values are indicated by color and square size. P values are indicated by white asterisks as * p <0.05, **p <0.01, *** p <0.001. T4 MBC and antibody data were preferred for Ad26.COV2.S due to fewer T5 paired samples. **(K-N)** The association of indicated parameters shown by scatter plot. Red indicated mRNA, green indicated Ad26.COV2.S. Spearman rank-order correlation values (r) and p values were shown. **(O)** Principal component analysis (PCA) representation of mRNA-1273 (n=19), BNT162b2 (n=14), Ad26.COV2.S (n=14), and NVX-Cov-2373 (n=10) on the basis of all parameters obtained 6-month post-vaccination. Only paired subjects were used for the PCA analysis. Arrows indicated the prominent immunological distinguishing features. Ellipse represented the clustering of each vaccine. Red indicated mRANA-1273, blue indicated BNT162b2, and green indicated Ad26.COV2.S. MBCs indicates spike-specific memory B cell, cMBCs indicates spike-specific classical memory B cell, aMBCs indicates spike-specific activated memory B cell, AIM1 + indicates OX40+CD137+, AIM2+ indicates OX40+CD40L+, nAb indicates neutralization antibody. **(P)** Spearman rank-order correlation between PSV neutralization titers and frequency of spike MBCs expressing CXCR3 at 3.5 months after vaccination. Background-subtracted and log data analyzed.

Next, we tested for relationships between early immune responses and immune memory (**Figures 7J-N, S13B** and **S14**). Peak post-2^nd^ mRNA immunization cTfh CD4^+^ T cells were strongly associated with 6-month antibody levels (**Figures 7J-L,** and **S13C-D**), providing an early indicator of long term humoral immunity. Early RBD IgG titers after the 1^st^ mRNA immunization were positively associated with 6-month RBD-specific memory B cell frequencies (**Figure S14E**-**F**). For both mRNA and Ad26.COV2.S, peak ICS^+^ CD4^+^ and CD8^+^ T cell responses significantly cross-correlated (T3, **Figure 7J**). Overall, these observations suggest that early peak CD4^+^ T cells responses had a lasting effect on the humoral response.

PCA mapping was performed using 3.5-month (**Figure S14G**) and 6-month (**Figure 7O**) post-vaccination data. PCA discriminated mRNA-1273 and Ad26.COV2.S, indicating these two vaccines generated distinct immunological profiles (**Figure 7O**). BNT162b2 largely developed the same profile as mRNA-1273 but with more heterogeneity. NVX-CoV2373 generated an immune memory profile overlapping with that of mRNA and adenoviral vectors (**Figure 7O**). Prominent immunological features distinguishing between mRNA and Ad26.COV2.S were CXCR3^+^ spike-specific memory B cells, ICS^+^ memory CD4^+^ T cells, CD71^+^ memory B cells, and spike IgG (**Figure 7O** and **S14G**). Notably, neutralizing antibody titers and CXCR3^+^ spike-specific memory B cells were correlated for Ad26.COV2.S vaccinees (r=0.44, p=0.04) but not mRNA vaccinees (mRNA-1273, p=0.25. BNT162b2, p=0.79. **Figure 7P**), corroborating the immunologically distinct outcomes. Overall, substantial relationships were observed between multiple components of immune memory for these COVID-19 vaccines, with distinct immune memory profiles for different vaccine platforms.

## DISCUSSION

COVID-19 vaccines have achieved extraordinary success in protection from infection and disease; yet some limitations exist, including differences in VE between vaccines and waning of protection against infection over a period of several months. Here, diverse metrics of adaptive responses were measured to mRNA-1273, BNT162b2, Ad26.COV2.S and NVX-CoV2373, with implications for understanding the protection against COVID-19 associated with each of the vaccines. A strength of this study is that the samples from different vaccine platforms were obtained from the same blood processing facility, from the same geographical location, and were analyzed concomitantly, utilizing the same experimental platform.

In the present study, antibody responses were detected in 100% of individuals. At 6 months post-immunization, the neutralizing antibody titer hierarchy between the vaccines was mRNA-1273~BNT162b2~NVX-CoV2373>Ad26.COV2.S. These serological data are consistent with previous reports for single vaccines (Atmar et al., 2022; Doria-Rose et al., 2021; Goel et al., 2021; Naranbhai et al., 2021a; Pajon et al., 2022; Pegu et al., 2021), and serological comparisons between vaccines (Barouch et al., 2021; Carreno et al., 2022; Naranbhai et al., 2021b; Self et al., 2021), though in much large serological studies ~2-fold higher neutralizing antibody titers were discerned with mRNA-1273 compared to BNT162b2 (Atmar *et al*., 2022; Steensels et al., 2021). Comparisons of NVX-CoV2373 antibody responses compared to other vaccines after 6 months have been very limited. Here we observed that NVX-CoV2373 neutralizing antibody titers were comparable to that of BNT162b2 and only moderately lower than mRNA-1273.

In this side-by-side comparative study, spike-specific CD4^+^ T cell responses were detected in 100% of individuals to all four vaccines. While neutralizing antibody kinetics were different between mRNA and viral vector vaccines, the CD4^+^ T cell response kinetics were similar. The hierarchy of the magnitude of the memory CD4^+^ T cells was mRNA-1273>BNT162b2~NVX-CoV2373>Ad26.COV2.S. These overall findings are consistent with previous reports on COVID-19 vaccine T cell responses (Barouch *et al*., 2021; Goel *et al*., 2021; Guerrera et al., 2021; Khoo et al., 2022; Liu et al., 2022; Mateus *et al*., 2021; Rodda et al., 2022; Tarke et al., 2022), but the analysis reported herein extensively expand these observations, including four different vaccines representing three different vaccine platforms, and with longitudinal data and single-cell cytokine expression resolution providing insights regarding CD4^+^ T cell subpopulations between the vaccines. Interestingly, multifunctional CD4^+^ T cells were observed most frequently after mRNA-1273 immunization, and CD4-CTLs represented a substantial fraction of the memory CD4^+^ T cells after mRNA-1273, BNT162b2, or NVX-CoV2373 vaccination. cTfh memory cells were represented as a substantial fraction of CD4^+^ T cell memory for each of the 4 vaccines, consistent with these vaccine platforms being selected for their ability to induce antibody responses. Memory CD4^+^ T cell responses were also comparted to infected individuals, demonstrated that each vaccine was successful in generating circulating spike-specific CD4^+^ T cell memory frequencies similar to or higher than SARS-CoV-2 infection, though of course infection also generates responses to other viral antigens.

The two mRNA vaccines and Ad26.COV2.S induced comparable acute and memory CD8^+^ T cell frequencies. These data are broadly consistent with previous reports for mRNA vaccines or adenoviral vectors (Goel *et al*., 2021; Guerrera *et al*., 2021; Keeton et al., 2022; Mateus *et al*., 2021; Tarke *et al*.,2022), with the exception being represented by reduced cytokine-expressing CD8^+^ T cells detected after mRNA vaccinations when using a 6 to 8-hr assay (Atmar *et al*., 2022; Collier et al., 2021), compared to the overnight stimulation used here. As expected for a protein-based vaccine, CD8^+^ T cell cytokine responses to NVX-CoV2373 were lower than all other vaccine platforms assessed, but it was notable that NVX-CoV2373 generated spike-specific CD8^+^ T cell memory in a fraction of individuals.

Spike- and RBD-specific memory B cell responses were detected in all individuals to each of the four vaccines. While neutralizing antibody titers declined over time in mRNA vaccinees, the frequency of spike-specific memory B cells increased over time. These divergent antibody and memory B cell kinetics were also observed in SARS-CoV-2 infection (Dan et al., 2021). The mRNA vaccine data are comparable to Goel et al. (Goel *et al*., 2021), but memory B cell data and kinetics for the Ad26.COV2.S or NVX-CoV2373 vaccines have not previously been available. At 6 months post-immunization, the spike-specific memory B cell hierarchy was mRNA1273~BNT162b2> Ad26.COV2.S>NVX-CoV2373. One of the most differentiating features of Ad26.COV2.S immunization observed here was the high frequency of CXCR3^+^ memory B cells. CXCR3^+^ memory B cells were correlated with neutralizing antibody titers after Ad26.COV2.S immunization, but not mRNA immunization, suggesting a specific functional role in viral vector B cell responses. CXCR3 expression on memory B cells has been found to be important for mucosal immunity in two mouse models (Oh et al., 2019; Oh et al., 2021).

Across the antigen-specific immune metrics assessed, mRNA vaccines were consistently the most immunogenic, with levels higher than or equal to that of Ad26.COV2.S and NVX-CoV2373 vaccines for each immune response. NVX-CoV2373 elicited CD4^+^ T cell memory and neutralizing antibody titers comparably to the mRNA vaccines. The responses induced by the Ad26.COV2.S were generally lower but relatively stable. The mRNA vaccine platforms were associated with substantial declines in neutralizing antibody titers over 6 months, while memory CD4^+^ T cells, memory CD8^+^ T cells, and memory B cells exhibited small reductions (T cells) or increases (B cells). These observations appear to be consistent with the relatively high degree of protection maintained against hospitalizations with COVID-19 after these vaccines over 6 months, and the differential VE reported between mRNA COVID-19 vaccines and Ad26.COV2.S. These results of detailed immunological evaluations, coupled with analyses of VE data published for the various vaccine platforms, may also be relevant for other vaccine efforts.

### Limitations of the Study

We did not evaluate recognition of variants, as this was evaluated in independent studies from our laboratories and others (Flemming, 2022; Gao et al., 2022; GeurtsvanKessel et al., 2022; Keeton et al., 2021; Tarke *et al*., 2022; Tarke et al., 2021). The current study did not evaluate responses elicited by other vaccine platforms (AstraZeneca, Coronavac, Sinopharm, Sputnik) commonly utilized in other regions because samples from individuals vaccinated with these platforms were not available to us.

## ACKNOWLEDGEMENTS

We thank Gina Levi and the LJI clinical core for assistance in sample coordination and blood processing. This project has been funded in whole or in part with federal funds from the National Institute of Allergy and Infectious Diseases, National Institutes of Health, Department of Health and Human Services, under grant CCHI AI142742 (S.C., A.S), Contract No. 75N9301900065 to A.S, D.W and U01 CA260541-01 (D.W). This work was additionally supported in part by LJI Institutional Funds and the NIAID under K08 award AI135078 (J.M.D.).

## AUTHOR CONTRIBUTIONS

Conceptualization: DW, AS, SC; Methodology: ZZ, JM, JMD, CHC, CRM, JC, EAE, BG, NB, DW; Formal analysis: JM, JMD, ZZ, CHC, CRM, JC, EAE, BG, NB, DW; Investigation: ZZ, JM, JMD, CHC, CRM, JC, EAE, BG, NB, DW; Project administration A.F., AS, SC, DW; Funding acquisition: JMD, AS, SC, DW; Writing: ZZ, JM, CHC, DW AS, SC; Supervision: DW, AS, SC.

## Key resources table

| REAGENT or RESOURCE | SOURCE | IDENTIFIER |
| --- | --- | --- |
| Antibodies |  |  |
| 11A9 (BUV496) [anti-CCR6] | BD Biosciences | Cat# 356920;<br>RRID: AB_2833076 |
| J252D4 (BV421) [anti-CXCR5] | BioLegend | Cat# 356920;<br>RRID: AB_2562303 |
| G025H7 (BV605) [anti-CXCR3] | BioLegend | Cat# 353728;<br>RRID: AB_2563157 |
| G043H7 (BV711) [anti-CCR7] | BioLegend | Cat# 353228;<br>RRID: AB_2563865 |
| G043H7 (PE/Cyanine7) [anti-CCR7] | BioLegend | Cat# 353226;<br>RRID: AB_11126145 |
| HB14 [anti-CD40] | Miltenyi Biotec | Cat# 130-108-041 |
| UCHT1 (BUV395) [anti-CD3] | BD Biosciences | Cat# 563546;<br>RRID: AB_2744387 |
| SK7 (PerCP) [anti-CD3] | BioLegend | Cat# 344814;<br>RRID: AB_10639948 |
| SK1 (BUV805) [anti-CD8] | BD Biosciences | Cat# 612889;<br>RRID: AB_2833078 |
| 3G8 (BV510) [anti-CD16] | BioLegend | Cat# 302048;<br>RRID: AB_2562085 |
| 3G8 (PerCP) [anti-CD16] | BioLegend | Cat# 302030;<br>RRID: AB_940380 |
| 63D3 (BV510) [anti-CD14] | BioLegend | Cat# 367124;<br>RRID: AB_2716229 |
| 63D3 (PerCP) [anti-CD14] | BioLegend | Cat# 367152 |
| 2H7 (BV510) [anti-CD20] | BioLegend | Cat# 302340;<br>RRID: AB_2561941 |
| HI100 (BV570) [anti-CD45RA] | BioLegend | Cat# 304132;<br>RRID: AB_2563813 |
| SK3 (cFluor548) [anti-CD4] | Cytek | Cat# R7-20043 |
| G46-6 (APC-R700) [anti-HLA-DR] | BD Biosciences | Cat# 565127;<br>RRID: AB_2732055 |
| DX2 (BB700) [anti-CD95] | BD Biosciences | Cat# 566542;<br>RRID: AB_2869780 |
| DX2 (BUV737) [anti-CD95] | BD Biosciences | Cat# 612790;<br>RRID: AB_2870117 |
| HB-7 (BV650) [anti-CD38] | BioLegend | Cat# 356620;<br>RRID: AB_2566233 |
| EH12.2H7 (BV785) [anti-PD-1] | BioLegend | Cat# 329930;<br>RRID: AB_2563443 |
| FN50 (FITC) [anti-CD69] | BioLegend | Cat# 310904;<br>RRID: AB_314839 |
| 24-31 (PE/Dazzle594) [anti-CD154/CD40L] | BioLegend | Cat# 310840;<br>RRID: AB_2566245 |
| 24-31 (PerCP-eFluor710) [anti-CD154/CD40L] | Thermo Fisher Scientific | Cat# 6-1548-42;<br>RRID: AB_10670357 |
| 4b4-1 (BUV737) [anti-CD137] | BD Bioscience | Cat# 741861;<br>RRID: AB_2871191 |
| Ber-Act35 (APC) [anti-CD134/OX40] | BioLegend | Cat# 350008;<br>RRID: AB_10719958 |
| 4S.B3 (FITC) [anti-IFNγ] | eBioscience | Cat# 11-7319-82;<br>RRID: AB_465415 |
| MP4-25D2 (BUV737) [anti-IL-4] | BD Bioscience | Cat# 612835;<br>RRID: AB_2870157 |
| BL168 (BV785) [anti-IL17] | BioLegend | Cat# 512338;<br>RRID: AB_2566765 |
| MQ1-17H12 (BB700) [anti-IL-2] | BD Bioscience | Cat# 566405;<br>RRID: AB_2744488 |
| JES3-19F1 (PE/Dazzle594) [anti-IL10] | BioLegend | Cat# 506812;<br>RRID: AB_2632783 |
| Mab11 (eFluor450) [anti-TNFα] | Thermo Fisher Scientific | Cat# 48-7349-42;<br>RRID: AB_2043889 |
| GB11 (Alexa Fluor 647) [anti-Granzyme B] | BD Bioscience | Cat# 560212 |
| Hu Fc Block Pure Fc1.3216 | BD Bioscience | Cat# 564220;<br>RRID: AB_2869554 |
| SJ25C1 (BUV563) [anti-CD19] | BD Biosciences | Cat# #612916;<br>RRID: AB_2870201 |
| 1C6 (BUV805) [anti-CXCR3] | BD Biosciences | Cat# 612790;<br>RRID: AB_2871338 |
| IA6-2 (Pacific Blue) [anti-IgD] | BioLegend | Cat# 348224 |
| MHM-88 (BV570) [anti-IgM] | BioLegend | Cat# 314517;<br>RRID: AB_10913816) |
| M-T271 (BB515) [anti-CD27] | BD Biosciences | Cat# 564642;<br>RRID: AB_2744354 |
| IS11-8E10 (Vio Bright) [anti-IgA] | Miltenyi Biotec | Cat# 130-113-480 |
| HCD56 (PerCP) [anti-CD56] | BioLegend | Cat# 318342;<br>RRID: AB_2561865 |
| M1310G05 (PerCP/Cyanine5.5) [anti-IgG] | BioLegend | Cat# 410710;<br>RRID: AB_2565788 |
| CY1G4 (PE/Dazzle594) [anti-CD71] | BioLegend | Cat# 334120;<br>RRID: AB_2734335 |
| 3.9 (PE/Cyanine5) [anti CD11c] | BioLegend | Cat# 301610;<br>RRID: AB_493578 |
| Bu31 (Alexa Fluor 700) [anti-CD21] | BioLegend | Cat# 354918;<br>RRID: AB_2750239 |
| HIT2 (APC/Fire810) [anti-CD38] | BioLegend | Cat# 303550;<br>RRID: AB_2860784 |
| HP6043 (Peroxidase) [anti-IgG] | Hybridoma Reagent Lab | Cat# HP6043-HRP |
| Biological samples |  |  |
| COVID-19 vaccinee donor blood samples | LJI Clinical Core | N/A |
| Chemicals, peptides, and recombinant proteins |  |  |
| Fixable Live/Dead Blue | Thermo Fischer Scientific | Cat# L34962 |
| SARS-CoV-2 Spike MP | Grifoni, 2020 | N/A |
| Brilliant Staining Buffer Plus | BD Biosciences | Cat# 566385<br>RRID: AB_2869761 |
| Brilliant Stain Buffer | BD Biosciences | Cat# 566349<br>RRID: AB_2869750 |
| SARS-CoV-2 Spike protein | Acro Biosystems | Cat# SPN-C82E9 |
| SARS-CoV-2 Spike protein Receptor-Binding Domain (RBD) | BioLegend | Cat# 793906 |
| Alexa Fluor 647 Streptavidin | Thermo Fisher Scientific | Cat# S21374 |
| BV421 Streptavidin | BioLegend | Cat# 405225 |
| BV711 Streptavidin | BioLegend | Cat# 405241 |
| PE-Cy7 Streptavidin | BioLegend | Cat# 405206 |
| PE-Cy5.5 Streptavidin | Thermo Fisher Scientific | Cat# SA1018 |
| Biotin | Avidity | Cat# Bir500A |
| Bacterial and virus strains |  |  |
| rVSV-SARS-CoV-2 | This study | N/A |
| Experimental models: Cell lines |  |  |
| VERO cells | ATCC | ATCC CCL-81 |
| HEK293T cells | ATCC | ATCC CRL-3216 |
| Software and algorithms |  |  |
| Flowjo 10.8.1 | FlowJo, LLC | www.flowjo.com |
| GraphPad Prism 9.3.0 | GraphPad | www.graphpad.com |
| Other |  |  |

## RESOURCE AVAILABILITY

### Lead contact

Further information and requests for resources and reagents should be directed to and will be fulfilled by the lead contact, Alessandro Sette

### Materials availability

Upon specific request and execution of a material transfer agreement (MTA) to the Lead Contact or to Daniela Weiskopf, aliquots of the peptide pools utilized in this study will be made available. Limitations might be applied to the availability of peptide reagents due to cost, quantity, demand, and availability.

### Data and code availability

All the data generated in this study are available in the published article and summarized in the corresponding tables, figures and supplemental materials.

## EXPERIMENTAL MODEL AND SUBJECT DETAILS

### Human Sample donors

A total of 354 peripheral blood samples were obtained from 102 participants who received either the Moderna mRNA-1273 (n=30), Pfizer/BioNTech BNT162b2 (n=30), Janssen Ad26.COV2.S (n=30) and Novavax NVX-CoV2373 vaccines (n=12), according to the approved dose schedule.

For baseline determinations, for a subset of donors, blood samples were collected before vaccination (T1), and subsequently 2 weeks (15 ± 3 days) after the first immunization (T2), 2 weeks (45 ± 35 days after first immunization) after the second immunization, 3.5 months (105 ± 7 days) and 6 months thereafter (185 ± 6 days). Both cohorts of mRNA vaccinees (mRNA-1273, Pfizer/BioNTech BNT162b2) received two doses of the vaccine (28 and 21 days apart, respectively). In the case of Ad26.COV2.S blood donations were collected after one dose at the same timepoints. Finally, in the case of the Novavax NVX-CoV2373, we advertised locally to recruit subjects who had participated in an investigational NVX-CoV2373 trial conducted in the San Diego region, where two intramuscular 5-μg doses of NVX-CoV2373 or placebo were administered 21 days apart (Clinicaltrials.gov). The study was structured in such way that donors received either first a placebo injection followed 21 later from the vaccine, or a vaccine injection followed by a placebo injection 21 days later; participants were blinded to their immunization regimen, and LJI had no information on which group the participants were in. An overview of samples analyzed in this study is provided in **Figure 1**. All experiments performed at the La Jolla Institute (LJI) were approved by the institutional review boards (IRB) of the La Jolla Institute (IRB#: VD-214).

To compare levels of immune memory responses induced by any of the vaccine platforms to immune memory responses induced by infection with SARS-CoV-2, samples were used from individuals that experienced infection with SARS-CoV-2, originally reported in (Mateus *et al*., 2021). We matched the 7 months (209 days) post-vaccination samples with samples from convalescent donors collected on average 181 days (range 170-195) post symptoms onset (PSOB cell experiments were repeated for 9 donors of this cohort and new 5 donors. The 5 new donors were selected randomly based to match the timepoint post symptom onset of the other samples. Seropositivity against SARS-CoV-2 was confirmed by ELISA, as described below. At the time of enrollment, all COVID-19 convalescent donors provided informed consent to participate in the present and future studies.

### Exclusion criteria

Before analyzing the entire data set for our cohort, we generated exclusion criteria as follows: subjects who tested positive for RBD and neutralization antibodies at baseline were excluded (one subject, mRNA-1273); subjects with no baseline sample available and whose RBD and neutralization antibody reached the peak after first-dose immunization (indicative of memory from previous infection) and were nucleocapsid (NC) antibody-positive were also excluded as previously infected subjects (one subject, mRNA-1273). In addition, any time points following a confirmed COVID-19 booster immunization were excluded for any subject (five subjects, BNT162b2, two Ad26.COV2.S, two NVX-CoV2373).

## METHOD DETAILS

### Peripheral blood mononuclear cells (PBMCs) and plasma isolation

Whole blood samples from subjects vaccinated with the mRNA-1273, BNT162b2, Ad26.COV2.S, or NVX-CoV2373 COVID-19 vaccine and convalescent samples after COVID-19 infection were collected at La Jolla Institute in heparin-coated blood bags and centrifuged for 15 min at 803 g to separate the cellular fraction and plasma. Blood samples were collected at the times described above. The plasma was then carefully removed from the cell pellet and stored at minus 20°C. PBMCs were isolated by density-gradient sedimentation using Ficoll-Paque (Lymphoprep, Nycomed Pharma, Oslo, Norway) as previously described (Dan *et al*., 2021; Grifoni et al., 2020; Mateus *et al*., 2021; Mateus et al., 2020; Rydyznski Moderbacher et al., 2020). Isolated PBMCs were cryopreserved in cell recovery media containing 10% Dimethyl sulfoxide (DMSO) (Gibco), supplemented with 10% heat-inactivated fetal bovine serum (FBS; Hyclone Laboratories, Logan UT), and stored in liquid nitrogen until used in the assays. Plasma samples were used for antibody measurements by ELISA and PSV neutralization assay and PBMC samples were used for flow cytometry in the T cell and B cell assays.

### SARS-CoV-2 ELISAs

The SARS-CoV-2 ELISAs have been described previously (Dan *et al*., 2021; Grifoni *et al*., 2020; Mateus *et al*., 2021; Rydyznski Moderbacher *et al*., 2020). Briefly, 96-well half-area plates (ThermoFisher 3690) were coated with 1 *u*g/mL of antigen and incubated at 4°C overnight. Antigens included recombinant SARS-CoV-2 RBD protein and spike protein, both obtained from the Saphire laboratory at LJI, and recombinant nucleocapsid protein (GenScript Z03488). The next day, plates were blocked with 3% milk in phosphate-buffered saline (PBS) containing 0.05% Tween-20 for 1.5 hours at room temperature. Plasma was heat-inactivated at 56°C for 30 to 60 min. Plasma was diluted in 1% milk containing 0.05% Tween-20 in PBS starting at a 1:3 dilution followed by serial dilutions by three and incubated for 1.5 hours at room temperature. Plates were washed five times with 0.05% PBS-Tween-20. Secondary antibodies were diluted in 1% milk containing 0.05% Tween-20 in PBS. Anti-human IgG peroxidase antibody produced in goat (Sigma A6029) was used at a 1:5,000 dilution. Plates were read on Spectramax Plate Reader at 450 nm, and data analysis was performed using SoftMax Pro.

End-point titers were plotted for each sample, using background-subtracted data. Negative and positive controls were used to standardize each assay and normalize across experiments. A positive control standard was created by pooling plasma from 6 convalescent COVID-19 donors to normalize between experiments. The limit of detection (LOD) was defined as 1:3 of IgG. The limit of quantification (LOQ) for COVID-19 vaccinated individuals were established based on pre-vaccinated individuals (timepoint 1) and set as the titer at which 95% of pre-vaccinated samples (T1) fell below the dotted line (**Figures 2A-B**). Titers, LOD, and LOQ were calibrated to the WHO International Reference Panel for anti-SARS-CoV-2 spike, RBD, and nucleocapsid binding antibody units per milliliter (WHO BAU/mL). For Spike IgG, the LOD was 0.20 with a LOQ of 1.024 (**Figures 2A** and **2D**). For RBD IgG, the LOD was 0.83 with a LOQ of 7.12 (**Figures 2B** and **2E**). For NC IgG, the LOD was 0.68 with a LOQ of 30.48 (**Figure S1A**).

For comparison among mRNA-1273, BNT162b, and Ad26.COV2.S over the entire 6+ month time period, log_10_ transformed end-point titers (WHO BAU/mL) were used to generate area under the curve (AUC) for each donor (**Figures S1B-D**). Donors with only 1 timepoint excluded. If there was no (T1), T1 was set as the LOD ET (BAU/mL). Correction factors for AUCs were determined by the number of time points and normalized to compare donor to donor. For comparison among mRNA-1273, BNT162b, Ad26.COV2.S, and NVZ-CoV2373 over the 3.5 months to 6 months period, log_10_ transformed end-point titers (WHO BAU/mL) were used to generate area under the curve (AUC) for each donor (**Figures S1E-G**). Donors with only 1 timepoint excluded. Kruskal-Wallis tests for AUC were <0.0001 for **Figures S1B-G**. Comparison between different vaccines were made by Mann-Whitney. Values plotted show GMT with GM SD.

### Pseudovirus (PSV) Neutralization Assay

The PSV neutralization assays in samples from vaccinated subjects were performed as previously described (Dan *et al*., 2021; Grifoni *et al*., 2020; Mateus *et al*., 2021; Rydyznski Moderbacher *et al*., 2020). Briefly, 2.5×10^4^ VERO cells (ATCC, Cat. No. CCL-81) were seeded in clear flat-bottom 96-well plates (Thermo Scientific, Cat. No. 165305) to produce a monolayer at the time of infection. Recombinant SARS-CoV-2-S-D614G pseudotyped VSV-ΔG-GFP were generated by transfecting HEK293T cells (ATCC, Cat. No. CRL-321) with plasmid phCMV3-SARS-CoV2-Spike kindly provided by Dr. E. Saphire and then infecting with VSV-ΔG-GFP. Pre-titrated rVSV-SARS-CoV-2-S-D614G was incubated with serially diluted human heat-inactivated plasma at 37°C for 1-1.5 hours before addition to confluent VERO cell monolayers. Cells were incubated for 16 hours at 37°C in 5% CO_2_ then fixed in 4% paraformaldehyde in PBS pH 7.4 (Santa Cruz, Cat. No. sc-281692) with 10 μg/ml of Hoechst (Thermo Scientific, Cat. No. 62249), and imaged using a CellInsight CX5 imager to quantify the total number of cells and infected GFP expressing cells to determine the percentage of infection. Neutralization titers or inhibition dose 50 (ID50) were calculated using the One-Site Fit Log IC50 model in Prism 9.3 (GraphPad). As internal quality control to define the variation inter-assay, a pooled plasma (secondary standard) from 10 donors who received the mRNA-1273 vaccine was included across the PSV neutralization assays. Samples that did not reach 50% inhibition at the lowest serum dilution of 1:20 were considered as non-neutralizing and the values were set to 19. PSV neutralization titers were done with two replicates per experiment. We included the WHO International Reference Panel for anti-SARS-CoV-2 immunoglobulin (20/268) to calibrate our PSV neutralization titers. The WHO IU calibrated neutralization ID50 (cID50-IU/mL) was graphed in figures. The limit of detection was calculated as 10.73 IU/mL.

### Spike megapool (Spike MP)

We have previously developed the MP approach to allow simultaneous testing of a large number of epitopes, as reported previously (Grifoni *et al*., 2020; Mateus *et al*., 2020; Rydyznski Moderbacher *et al*.,2020). According to this approach, large numbers of different epitopes are solubilized, pooled, and relyophilized to avoid cell toxicity problems associated with high concentrations of DMSO typically encountered when single pre-solubilized epitopes are pooled (Grifoni *et al*., 2020; Mateus *et al*., 2020; Rydyznski Moderbacher *et al*., 2020). Here, were used for ex vivo stimulation of PBMCs for flow cytometry a MP to evaluate the antigen-specific T cell response against SARS-CoV-2 spike. We used a Spike MP of 253 overlapping peptides spanning the entire sequence of the Spike protein. As this peptide pool consists of peptides with a length of 15 amino acids, both CD4^+^ and CD8^+^ T cells have the capacity to recognize this MP, as described previously (Dan *et al*., 2021; Mateus *et al*., 2021).

### Activation-induced markers (AIM) assay

The AIM assays in samples from subjects vaccinated with mRNA-1273, BNT162b2, Ad26.COV2.S, or NVX-CoV2373 COVID-19 vaccine were performed as previously described (Dan *et al*., 2021; Grifoni *et al*., 2020; Mateus *et al*., 2021; Mateus *et al*., 2020; Rydyznski Moderbacher *et al*., 2020).

Spike-specific T cells were measured as a percentage of AIM^+^ (OX40^+^CD137^+^) CD4^+^ and (CD69^+^CD137^+^) CD8^+^ T cells after stimulation of PBMCs from subjects vaccinated with the Spike MP. Also, Spike-specific circulating T follicular helper (cTfh) cells (CXCR5^+^OX40^+^CD40L^+^, as a percentage of CD4^+^ T cells) were defined by the AIM assay. Briefly, prior to the addition of the Spike MP, PBMCs were blocked at 37°C for 15 min with 0.5 μg/ml anti-CD40 mAb (Miltenyi Biotec). Then, cells were incubated at 37°C for 24 hours in the presence of fluorescently labeled chemokine receptor antibodies (anti-CCR6, CXCR5, CXCR3, and CCR7) and the Spike MP (1 μg/ml) in 96-wells U-bottom plates, as previously described (Mateus *et al*., 2021; Rydyznski Moderbacher *et al*., 2020). In addition, PBMCs were incubated with an equimolar amount of DMSO as negative control and with phytohemagglutinin (5 μg/ml) (PHA, Roche) as a positive control. For the surface stain, 1×10^6^ PBMCs were resuspended in PBS, incubated with BD human FC block (BD Biosciences, San Diego, CA) and the LIVE/DEAD marker in the dark for 15 min and washed with PBS. Then, the antibody mix containing the rest of the surface antibodies was added directly to cells and incubated for 60 min at 4°C in the dark. Following surface staining, cells were then washed twice with PBS containing 3% FBS (FACS buffer). All samples were acquired on a Cytek Aurora (Cytek Biosciences, Fremont, CA). A list of antibodies used in this panel can be found in **table S1** and a representative gating strategy of Spike-specific CD4^+^ and CD8^+^ T cells using the AIM assay is shown in **Figure S2**, respectively.

Spike-specific CD4^+^ and CD8^+^ T cells were measured as background (DMSO) subtracted data, with a minimal DMSO level set to 0.005%. Response > 0.02% and a stimulation index (SI) > 2 for CD4^+^ and > 0.03% and SI > 3 for CD8^+^ T cells were considered positive. The limit of quantification (LOQ) for antigen-specific CD4^+^ T cell responses (0.03%) and antigen-specific CD8^+^ T cell responses (0.05%) was calculated using the median two-fold standard deviation of all negative controls.

### Intracellular cytokine staining (ICS) assay

The ICS assays in samples from subjects vaccinated with mRNA-1273, BNT162b2, Ad26.COV2.S, or NVX-CoV2373 COVID-19 vaccine were performed as previously described (Mateus *et al*., 2021; Rydyznski Moderbacher *et al*., 2020).

Prior to the addition of the Spike MP, PBMC were blocked at 37°C for 15 minutes with 0.5 μg/ml anti-CD40 mAb, as previously described (Mateus *et al*., 2021; Rydyznski Moderbacher *et al*., 2020). PBMCs were cultured in the presence of the Spike MP (1 μg/ml) for 24 hours at 37°C in 96-wells U-bottom plates. In addition, cells were incubated with an equimolar amount of DMSO as a negative control. After 24 hours, Golgi-Plug and Golgi-Stop were added to the culture for 4 hours along with the anti-CD69 Ab. Cells were then washed, incubated with BD human FC block, and stained with the LIVE/DEAD marker as described above. Then, cells were washed and surface stained for 30 min at 4°C in the dark and fixed with 1% of paraformaldehyde (Sigma-Aldrich, St. Louis, MO). Subsequently, cells were permeated and stained with intracellular antibodies for 30 min at room temperature in the dark. All samples were acquired on a Cytek Aurora. Antibodies used in the ICS assay are listed in **table S2** and a representative gating strategy of cytokine-producing spike-specific CD4^+^ and CD8^+^ T cells using the ICS assay is shown in **Figures 4A** and **5A**.

To define the spike-specific T cells by the ICS assay, we gated the cytokine- or GzB-producing cells together with the expression of iCD40L or CD69 on CD4^+^ or CD8^+^ T cells, respectively (**Figures 4A** and **5A**). Then, a Boolean analysis was performed to define the multifunctional profiles on FlowJo 10.8.1. The overall response to spike, denoted as Secreted-effector^+^ (IFNγ, TNFα, IL-2, and/or GzB) or Cytokine^+^ (IFNγ, TNFα, and/or IL-2), was defined as the sum of the background-subtracted responses to each combination of individual cytokines or GzB. The total spike-specific CD4^+^ and CD8^+^ T cells producing IFNγ, TNFα, IL-2, and/or GzB are shown in **Figures 4–5** and **Figure S5**, respectively. To define the multifunctional profiles of spike-specific T cells, all positive background-subtracted data (> 0.005% and a SI > 2 for CD4^+^ T cells and CD8^+^ T cells) was aggregated into a combined sum of antigen-specific CD4^+^ or CD8^+^ T cells based on the number of functions. Values higher than the LOQ (0.01%) were considered for the analysis of the multifunctional spike-specific T cell responses. The average of the relative CD4^+^ and CD8^+^ T cell responses was calculated per donor and visit to define the proportion of multifunctional spike-specific T cell responses with one, two, three, and four functions (**Figures 3E**, **5C**, **S4** and **S9**).

### Detection of SARS-CoV-2-specific memory B cells

Detection of SARS-CoV-2-specific memory B cells (MBCs) in samples from subjects vaccinated with mRNA-1273, BNT162b2, Ad26.COV2.S, or NVX-CoV2373 COVID-19 vaccine was performed using B cell probes as previously described (Dan *et al*., 2021) Biotinylated full-length SARS-CoV-2 Spike protein was purchased from Acro Biosystems and SARS-CoV-2 Spike protein Receptor-Binding Domain (RBD) was purchased from BioLegend.

To enhance specificity, identification of both spike- and RBD-specific MBCswas performed using two fluorochromes for each protein. Thus, the biotinylated SARS-CoV-2 spike was incubated with either Alexa Fluor 647 or BV421 at a 20:1 ratio (~6:1 molar ratio) for 1 hour at 4°C. Biotinylated RBD was conjugated with BV711 or PE-Cy7 at a 2.2:1 ratio (~4:1 molar ratio). Streptavidin PE-Cy5.5 was used as a decoy probe to minimize background by eliminating SARS-CoV-2 nonspecific streptavidin-binding B cells. Then, 9×10^6^ PBMCs were placed in U-bottom 96 well plates and stained with a solution consisting of 5 μM of biotin to avoid cross-reactivity among probes, 20 ng of decoy probe, 211 ng of Spike and 31.25 ng of RBD per sample, diluted in Brilliant Buffer and incubated for 1 hour at 4 °C, protected from light. After washing with PBS, cells were incubated with surface antibodies (**table S3**) diluted in Brilliant Staining Buffer for 30 min at 4°C in dark. Viability staining was performed using Live/Dead Fixable Blue Stain Kit diluted at 1:200 in PBS and incubation at 4°C for 30 minutes.

The acquisition was performed using Cytek Aurora. The frequency of antigen-specific MBCs was expressed as a percentage of total memory B cells (Singlets, Lymphocytes, Live, CD3– CD14– CD16– CD56–CD19+ CD20+ CD38int/– IgD– and/or CD27+). For every experiment, PBMCs from a known positive control (COVID-19 convalescent subject) and an unexposed subject were included to ensure consistent sensitivity and specificity of the assay. Limit of detection was calculated as median + 2x standard deviation (SD) of [1/(number of total B cells recorded)]

### Correlation and principal component analysis (PCA)

Correlograms plotting the Spearman rank correlation coefficient (r) between all paired parameters were created with the corrplot package (v0.84) running in Rstudio (1.1.456) as previously described (Rydyznski Moderbacher *et al*., 2020). Spearman rank two-tailed P values were calculated using corr.mtest and graphed based on * p < 0.05, **p < 0.01, *** p < 0.001. The codes used are:

M=cor(DataFrame, method=“spearman”, use = “pairwise.complete.obs”)
MP=cor.mtest(DataFrame, method=“spearman”, use = “pairwise.complete.obs”, conf.level=0.95, exact=FALSE)
corrplot(M, p.mat = MP$p, method = ‘square’, tl.col=“black”, tl.cex = 0.7, tl.srt = 45, cl.align=“l”, type = ‘lower’, sig.level = c(0.001, 0.01,0.05), pch.cex = 0.7, insig = ‘label_sig’, pch.col = ‘white’)
The “DataFrame” is the data from each correlation matrix shown in figure 6, collected and organized in spreadsheet.

The codes for PCA analysis are as follows:

res.pca=PCA(na.omit(MP), scale = TRUE)
fviz_eig(res.pca, addlabels = TRUE)
fviz_pca_biplot(res.pca, label =“var”, labelsize = 3, repel= TRUE, geom.ind = “point”, pointsize= 4, col.ind = Group$Vaccine, palette = c(“darkgreen”, “blue”, “red”, “purple”), col.var = “black”, alpha.var = 0.5, addEllipses = TRUE, ellipse.alpha=0, select.var=list(name=c(“RBD IgG”, “Spike IgG”, “nAbs”, “MBC”, “aMBC”, “cMBC”, “CXCR3+ MBC”, “AIM2+ CD4”, “ICS+ CD4”, “ICS+ CD8”)), ellipse.level=0.8, legend.title = “Groups”, invisible = “quali”, title = “”)

### Statistical analysis

Cytometry data was analyzed using FlowJo 10.8.1. Statistical analyses were performed in GraphPad Prism 9.3.0, unless otherwise stated. The statistical details of the experiments are provided in the respective figure legends. Data plotted in linear scale were expressed as Mean ±Standard Deviation (SD). Data plotted in logarithmic scales were expressed as Geometric Mean ±Geometric Standard Deviation (SD). Mann–Whitney U or Wilcoxon tests were applied for unpaired or paired comparisons, respectively. Kruskal– Wallis and Dunn’s posttest were also applied for multiple comparisons in vaccine cohorts. Details pertaining to significance are also noted in the respective legends.

