## Supplementary Materials for "Humoral and cellular immune memory to four COVID-19 vaccines"

### Supplemental Tables

**Table S1.** Reagents used for AIM assays.

| Reagent | Clone (Source) | Catalog. No. | Dilution |
|---|---|---|---|
| Live/Dead Blue | (ThermoFisher) | L23105 | 1:1000 |
| CCR6-BUV496 | 11A9 (BD Biosciences) | 612948 | 1:200 |
| CXCR5-BV421 | J252D4 (Biolegend) | 356920 | 1:200 |
| CXCR3-BV605 | G025H7 (Biolegend) | 353728 | 1:200 |
| CCR7-BV711 | G043H7 (Biolegend) | 353228 | 1:200 |
| CD3-BUV395 | UCHT1 (BD Biosciences) | 563546 | 1:1000 |
| CD137-BUV737 | 4B4-1 (BD Biosciences) | 741861 | 1:100 |
| CD8-BUV805 | SK1 (BD Biosciences) | 612889 | 1:1000 |
| CD16-BV510 | 3G8 (Biolegend) | 302048 | 1:1000 |
| CD14-BV510 | 63D3 (Biolegend) | 367124 | 1:1000 |
| CD20-BV510 | 2H7 (Biolegend) | 302340 | 1:1000 |
| CD45RA-BV570 | HI100 (Biolegend) | 304132 | 1:1000 |
| CD38-BV650 | HB-7 (Biolegend) | 356620 | 1:200 |
| PD-1-BV785 | EH12.2H7 (Biolegend) | 329930 | 1:200 |
| CD69-FITC | FN50 (Biolegend) | 310904 | 1:200 |
| CD4-cFluor b548 | SK3 (Cytek Biosciences) | R7-20043 | 1:500 |
| CD95-BB700 | DX2 (BD Biosciences) | 566542 | 1:500 |
| CD40L-PE-Dazzle594 | 24-31 (Biolegend) | 310840 | 1:200 |
| OX40-APC | Ber-Act35 (Biolegend) | 350008 | 1:100 |
| HLA-DR-APC-R700 | G46-6 (BD Biosciences) | 565127 | 1:500 |

**Table S2.** Reagents used for ICS assays.

| Reagent | Clone (Source) | Catalog. No. | Dilution |
|---|---|---|---|
| Live/Dead Blue | (ThermoFisher) | L23105 | 1:1000 |
| CD3-BUV395 | UCHT1 (BD Biosciences) | 563546 | 1:100 |
| CD8-BUV805 | SK1 (BD Biosciences) | 612889 | 1:100 |
| CD16-BV510 | 3G8 (Biolegend) | 302048 | 1:200 |
| CD14-BV510 | 63D3 (Biolegend) | 367124 | 1:200 |
| CD20-BV510 | 2H7 (Biolegend) | 302340 | 1:200 |
| CD45RA-BV570 | HI100 (Biolegend) | 304132 | 1:50 |
| CD4-cFluor b548 | SK3 (Cytek Biosciences) | R7-20043 | 1:25 |
| CD69-BV605 | FN50 (BD Biosciences) | 562989 | 1:100 |
| CCR7-PE-Cy7 | G043H7 (Biolegend) | 353226 | 1:100 |
| IL-4-BUV737 | MP4-25D2 (BD Biosciences) | 612835 | 1:200 |
| IL-17-BV785 | BL168 (Biolegend) | 512338 | 1:100 |
| IFNγ-FITC | 4S.B3 (ThermoFisher) | 11-7319-82 | 1:500 |
| IL-2-BB700 | MQ1-17H12 (BD Biosciences) | 566405 | 1:200 |
| IL-10 -PE-Dazzle594 | JES3-19F1 (Biolegend) | 506812 | 1:100 |
| TNFα-eFluor450 | Mab11 (ThermoFisher) | 48-7349-42 | 1:200 |
| Granzyme B-AF647 | GB11 (BD Biosciences) | 560212 | 1:50 |
| CD40L-PerCP-ef710 | 24-31 (ThermoFisher) | 46-1548-42 | 1:50 |

**Table S3.** Reagents used for B cell assays.

| Reagent | Clone (Source) | Catalog. No. | Dilution |
|---|---|---|---|
| Live/Dead Blue | (Thermo Fisher) | L34962 | 1:200 |
| CD19-BUV563 | SJ25C1 (BD Biosciences) | 612916 | 1:200 |
| CD95-BUV737 | DX2 (BD Biosciences) | 612790 | 1:200 |
| CD183-BUV805 | 1C6/CXCR3 (BD Biosciences) | 742048 | 1:50 |
| IgD-Pacific Blue | IA6-2 (Biolegend) | 348224 | 1:50 |
| CD20-Brilliant Violet 510 | 2H7 (Biolegend) | 302340 | 1:100 |
| IgM-Brilliant Violet 570 | MHM-88 (Biolegend) | 314517 | 1:200 |
| CD27-BB515 | M-T271 (BD Biosciences) | 564642 | 1:200 |
| IgA-VioBright-FITC | IS11-8E10 (Miltenyi Biotec) | 130-113-480 | 1:400 |
| CD3-PerCP | SK7 (Biolegend) | 344814 | 1:100 |
| CD14-PerCP | 63D3 (Biolegend) | 367152 | 1:200 |
| CD16-PerCP | 3G8 (Biolegend) | 302030 | 1:200 |
| CD56-PerCP | HCD56 (Biolegend) | 318342 | 1:200 |
| IgG-PerCP-Cyanine5.5 | M1310G05 (Biolegend) | 410710 | 1:100 |
| CD71-PE-Dazzle 594 | CY1G4 (Biolegend) | 334120 | 1:200 |
| CD11c-PE-Cy5 | 3.9 (Biolegend) | 301610 | 1:200 |
| CD21-Alexa Fluor 700 | Bu32 (Biolegend) | 354918 | 1:50 |
| CD38-APC-Fire 810 | HIT2 (Biolegend) | 303550 | 1:200 |
| Alexa Fluor 647 Streptavidin | (Thermo Fisher) | S21374 | 1:59 |
| BV421 Streptavidin | (Biolegend) | 405225 | 1:59 |
| BV711 Streptavidin | (Biolegend) | 405241 | 1:400 |
| PE-Cy7 Streptavidin | (Biolegend) | 405206 | 1:400 |
| PE-Cy5.5 Streptavidin | (Thermo Fisher) | SA1018 | 1:500 |

### Supplemental Figures

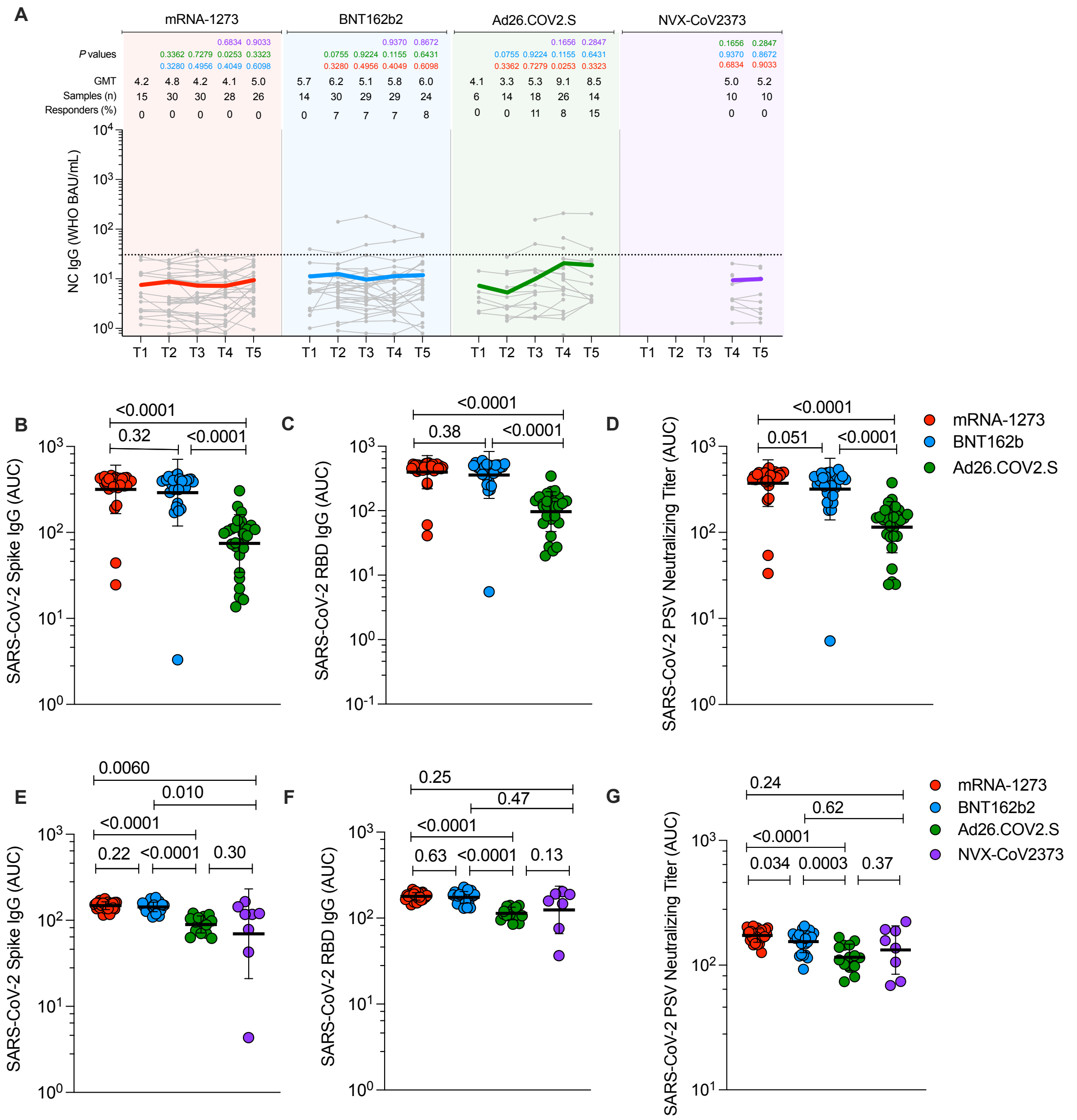


**Figure S1. Antibodies elicited by mRNA-1273, BNT162b2, Ad26.COV2.S, and NVX-CoV2373 COVID-19 vaccine platforms.**
**(A)** Comparison of longitudinal SARS-CoV-2 spike Nucleocapsid (NC) levels from all donors to the mRNA-1273 (red), BNT162b2 (blue), Ad26.COV2.S (green) and NVX-CoV2373 (purple) over 6 months. Individual subjects are show as gray symbols with connecting lines for longitudinal samples. Geometric means of overall responses are shown in thick colored lines. The dotted line indicates the limit of quantification (LOQ). LOQ was established on the basis of pre-vaccinated samples (timepoint 1) and set as the titer at which 95% of pre-vaccinated samples (T1) fell below the dotted line. P values on the top show the differences between each time point and vaccine between the different vaccines, color-coded per comparison based on the vaccine compared. NS, non-significant; GMT, Geometric mean titers.
**(B-D)** (B) Comparison of area under the curve (AUC) for spike IgG, (C) RBD IgG, and (D) PSV neutralization titers across the full 6-month window for mRNA-1273, BNT162b2, and Ad26.COV2.S. Statistical analysis by Mann-Whitney t-test.
**(E-G)** (E) Comparison of area under the curve (AUC) for spike IgG, (F) RBD IgG, and (G) PSV neutralization titers across the 3.5 to 6-month window for mRNA-1273, BNT162b2, Ad26.COV2.S, and NVX-COV2373. Statistical analysis by Mann-Whitney t-test.

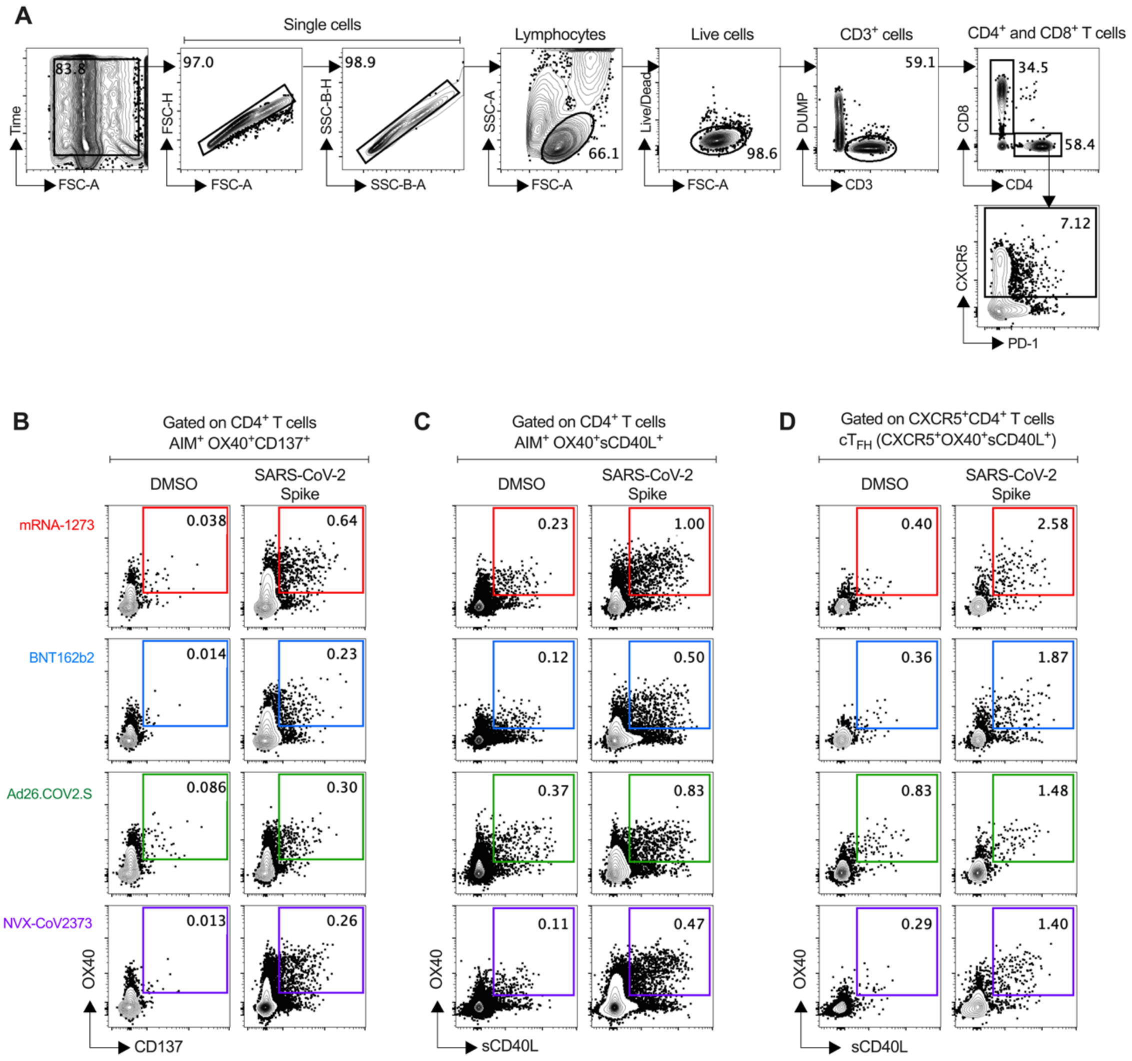


**Figure S2. Representative gating strategy for T cell analysis.**

**(A)** Representative strategies to define CD3$^+$CD4$^+$ and CD3$^+$CD8$^+$ cells by AIM and ICS assays.

**(B-C)** Representative gating strategy of spike-specific AIM$^+$ CD4$^+$ T cells induced by mRNA-1273, BNT162b2, Ad26.COV2.S, and NVX-CoV2373 COVID-19 vaccine platforms. Spike-specific CD4$^+$ T cells were measured by Activation-Induced Makers (AIM) assay: AIM$^+$ OX40$^+$ and CD137$^+$ (B) and AIM$^+$ OX40$^+$ and surface CD40L$^+$ (C). See **Figure S3** for AIM$^+$ OX40$^+$sCD40L results.

**(D)** Representative gating strategy of spike-specific circulating follicular helper T cells (c$T_{FH}$) induced by mRNA-1273, BNT162b2, Ad26.COV2.S, and NVX-CoV2373 COVID-19 vaccine platforms.

mRNA-1273 (red), BNT162b2 (blue), Ad26.COV2.S (green), and NVX-CoV2373 (purple).

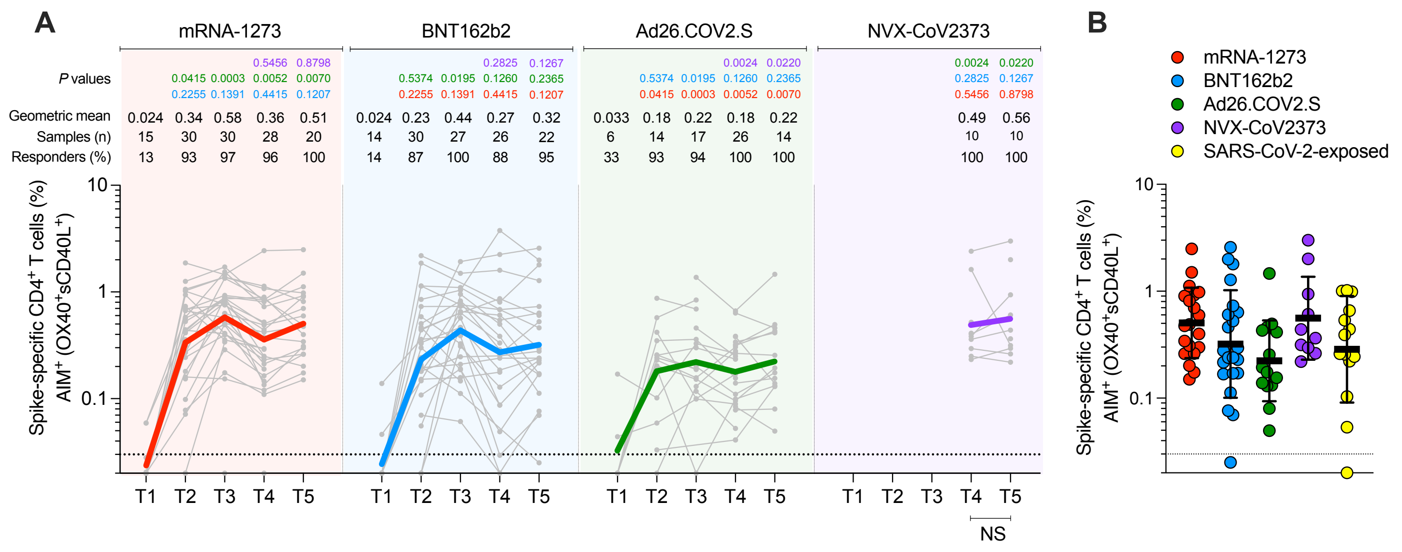


**Figure S3. Spike-specific CD4⁺ T cell responses induced by COVID-19 vaccines measured by AIM (OX40⁺sCD40L⁺).**

**(A)** Longitudinal spike-specific CD4⁺ AIM⁺ T cell responses induced by COVID-19 vaccines evaluated at T1, T2, T3, T4, and T5. Spike-specific CD4⁺ T cell frequencies were quantified by the expression of AIM surface markers (OX40⁺ sCD40L⁺) after stimulation with spike megapool (MP). See **Figure S2C** for the representative gating strategy of AIM⁺ cells.

**(B)** Comparison of spike-specific CD4⁺ T cell responses (AIM⁺ OX40⁺sCD40L⁺) between COVID-19 vaccinees at 185 ± 6 days post-vaccination and SARS-CoV-2-exposed subjects 170 to 195 days PSO.

The dotted black line indicates the limit of quantification (LOQ). The color-coded bold lines in (A) represent the Geometric mean in each time post-vaccination. Background-subtracted and log data analyzed. P values on the top in (A) show the differences between each time point in the different vaccines and are color-coded as follows: mRNA-1273 (red), BNT162b2 (blue), Ad26.COV2.S (green), or NVX-CoV2373 (purple). Data were analyzed for statistical significance using the Mann-Whitney test [(A-B)]. T1, Baseline; T2, 15 ± 3 days; T3, 42 ± 7 days; T4, 108 ± 9 days; T5, 185 ± 8 days; NS, non-significant.

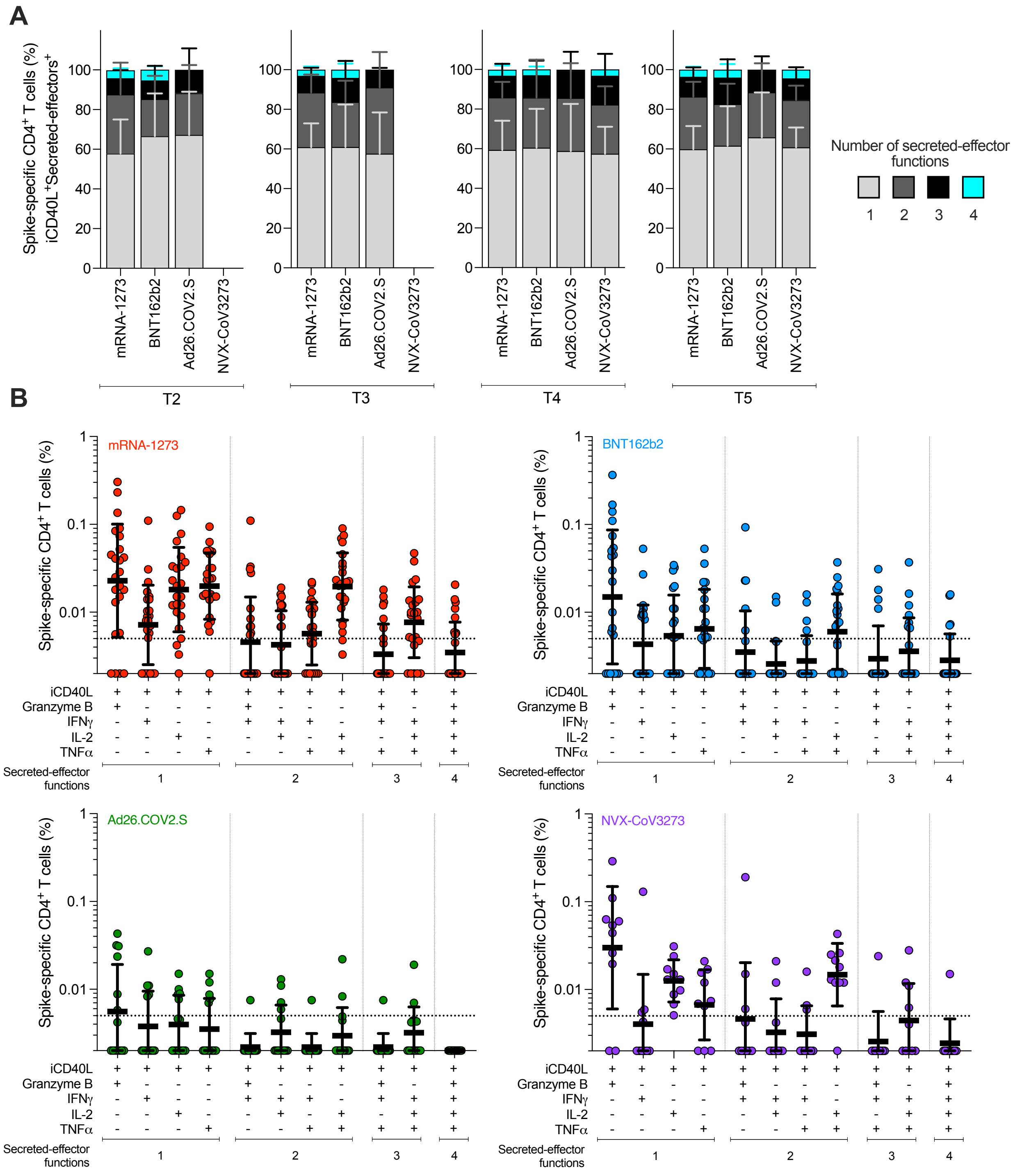


**Figure S4**. **Multifunctional spike-specific CD4⁺ T cells expressing iCD40L⁺ in subjects vaccinated with the mRNA-1273, BNT162b2, Ad26.COV2.S, or NVX-CoV2373 COVID-19 vaccines.**

**(A)** Comparison of multifunctional profiles of spike-specific CD4⁺ T cells iCD40L⁺Secreted-effector⁺ in subjects vaccinated with the mRNA-1273, BNT162b2, Ad26.COV2.S, or NVX-CoV2373 COVID-19 vaccine at T2, T3, T4, and T5. The blue, green, yellow, and red colors in the stacked bar charts depict the production of one, two, three, and four Secreted-effector⁺ functions, respectively. Data were analyzed for statistical significance using the Kruskal-Wallis (KW) test and Dunn's post-test for multiple comparisons.

**(B)** Predominant multifunctional profiles of spike-specific CD4⁺ T cells expressing iCD40L with one, two, three, and four Secreted-effector⁺ functions were analyzed in subjects vaccinated with the mRNA-1273, BNT162b2, Ad26.COV2.S, or NVX-CoV2373 COVID-19 vaccine at 6 months postvaccination (T5). Boolean analysis was carried out to define the functional profiles and the analysis included GzB, IFNγ, IL-2, and TNFα gated on CD3⁺CD4⁺ cells expressing iCD40L (See **Figure 4**). Each Secreted-effector⁺ profile combination was considered positive with >0.005% and a SI>2 for CD4⁺ T cells. The dotted line indicates the limit of quantification (LOQ). The bars show the Geometric mean and geometric SD of the spike-specific CD4⁺ T cells iCD40L⁺.

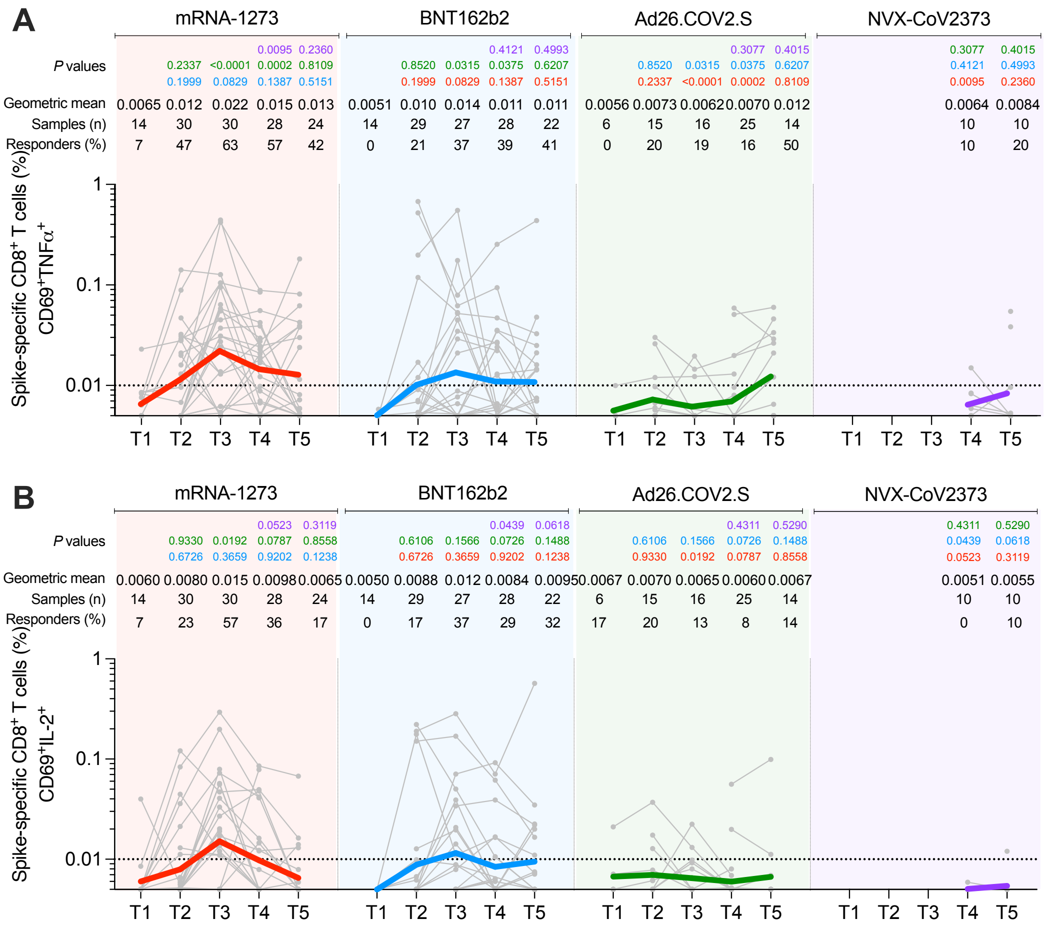

**Figure S5. Longitudinal spike-specific CD8⁺ T cells expressing intracellular CD69⁺ and producing cytokines in subjects vaccinated with the mRNA-1273, BNT162b2, Ad26.COV2.S, or NVX-CoV2373 COVID-19 vaccines.**

**(A-B)** Spike-specific $CD8^+$ T cells expressing $CD69^+$ and producing TNFα (A) and IL-2 (B) from COVID-19 vaccinees evaluated at T1, T2, T3, T4, and T5.

The dotted black line indicates the limit of quantification (LOQ). The color-coded bold lines in (A-B) represent the Geometric mean in each time post-vaccination. Background-subtracted and log data analyzed. P values on the top in (A-B) show the differences between each time point in the different vaccines and are color-coded as follows: mRNA-1273 (red), BNT162b2 (blue), Ad26.COV2.S (green), or NVX-CoV2373 (purple). Data were analyzed for statistical significance using the Mann-Whitney test [(A-B)]. T1, Baseline; T2, 15 ± 3 days; T3, 42 ± 7 days; T4, 108 ± 9 days; T5, 185 ± 8 days.

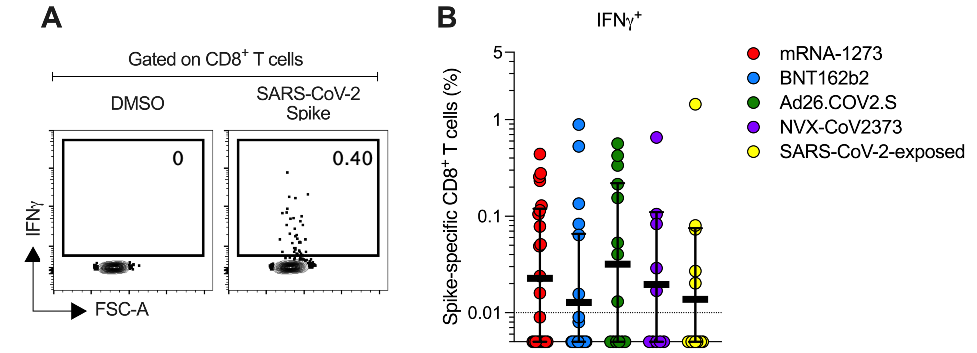


**Figure S6. Spike-specific CD8⁺ T cell responses producing IFNγ between COVID-19 vaccinees and SARS-CoV-2-exposed subjects.**

**(A)** Representative gating strategy of spike-specific CD8⁺ T cells producing IFNγ detected in COVID-19 vaccine platforms.

**(B)** Comparison of spike-specific CD8⁺ T cell responses producing IFNγ between COVID-19 vaccinees at 185 ± 6 days post-vaccination and SARS-CoV-2-exposed subjects 170 to 195 days PSO.

The dotted black line indicates the limit of quantification (LOQ). Background-subtracted and log data analyzed. T1, Baseline; T2, 15 ± 3 days; T3, 42 ± 7 days; T4, 108 ± 9 days; T5, 185 ± 8 days; NS, non-significant.

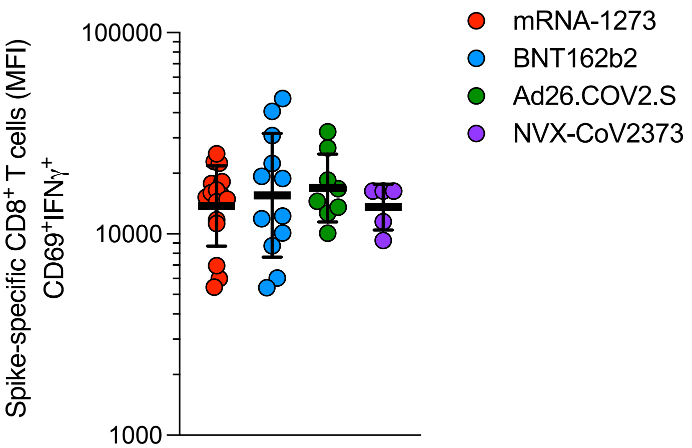


**Figure S7. Spike-specific CD8$^+$ T cells expressing CD69$^+$ and producing IFNγ from COVID-19 vaccinees evaluated at 6 months post-vaccination (T5).** Median fluorescence intensity (MFI) levels of IFNγ were evaluated on COVID-19 vaccinees with a positive IFNγ response detected at T5 (See **Figure 5**).

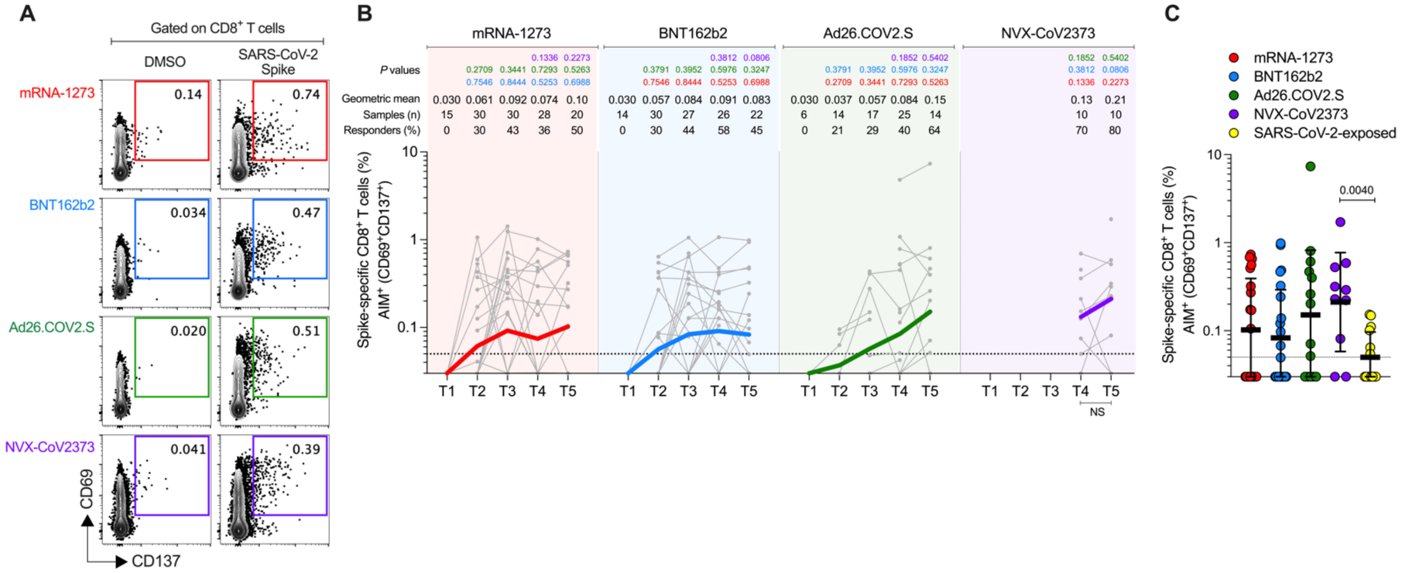

**Figure S8. Spike-specific CD8⁺ T cell responses induced by COVID-19 vaccines measured by AIM (CD69⁺CD137⁺).**

**(A)** Representative gating strategy of spike-specific AIM⁺ CD8⁺ T cells induced by mRNA-1273, BNT162b2, Ad26.COV2.S, and NVX-CoV2373 COVID-19 vaccine platforms. Spike-specific CD8⁺ T cells were measured by Activation-Induced Makers (AIM) assay: AIM⁺ CD69⁺ and CD137⁺ after stimulation with spike megapool (MP).

**(B)** Longitudinal spike-specific CD8⁺ AIM⁺ T cell responses induced by COVID-19 vaccines evaluated at T1, T2, T3, T4, and T5.

**(C)** Comparison of spike-specific CD8⁺ T cell responses (AIM⁺ CD69⁺CD137⁺) between COVID-19 vaccinees at 185 ± 6 days post-vaccination and SARS-CoV-2-exposed subjects 170 to 195 days PSO.

The dotted black line indicates the limit of quantification (LOQ). The color-coded bold lines in (B) represent the Geometric mean in each time post-vaccination. Background-subtracted and log data analyzed. P values on the top in (A) show the differences between each time point in the different vaccines and are color-coded as follows: mRNA-1273 (red), BNT162b2 (blue), Ad26.COV2.S (green), or NVX-CoV2373 (purple). Data were analyzed for statistical significance using the Mann-Whitney test [(B-C)]. T1, Baseline; T2, 15 ± 3 days; T3, 42 ± 7 days; T4, 108 ± 9 days; T5, 185 ± 8 days; NS, non-significant.

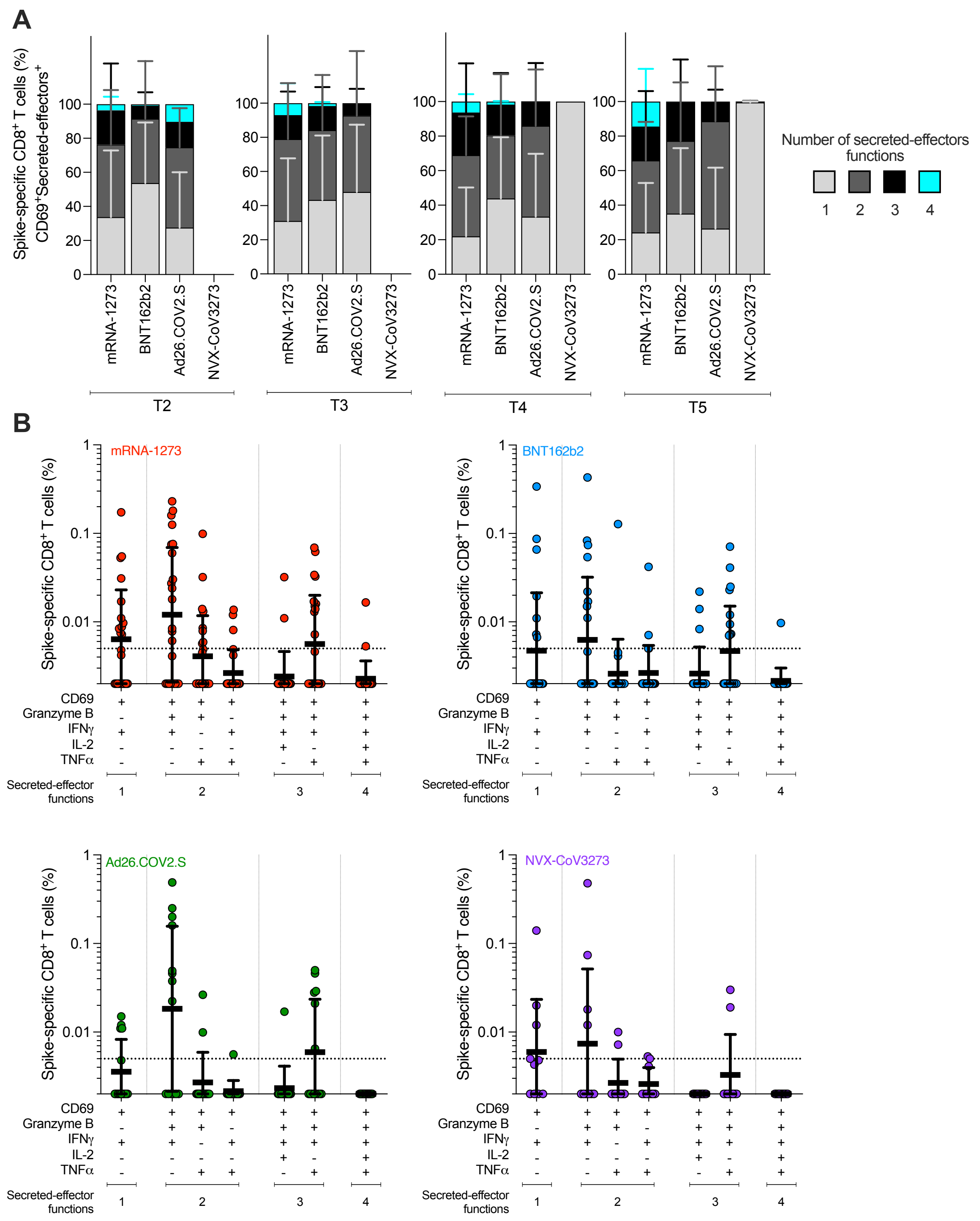


**Figure S9. Multifunctional spike-specific CD8⁺ T cells expressing CD69⁺ in subjects vaccinated with the mRNA-1273, BNT162b2, Ad26.COV2.S, or NVX-CoV2373 COVID-19 vaccines.**

**(A)** Comparison of multifunctional profiles of spike-specific CD8⁺ T cells CD69⁺Cytokine⁺ in subjects vaccinated with the mRNA-1273, BNT162b2, Ad26.COV2.S, or NVX-CoV2373 COVID-19 vaccine at T2, T3, T4, and T5. The blue, green, yellow, and red colors in the stacked bar charts depict the production of one, two, three, and four Cytokine⁺ functions, respectively. Data were analyzed for statistical significance using the Kruskal-Wallis (KW) test and Dunn's post-test for multiple comparisons.

**(B)** Predominant multifunctional profiles of spike-specific CD8⁺ T cells expressing CD69 with one, two, three, and four Cytokine⁺ functions were analyzed in subjects vaccinated with the mRNA-1273, BNT162b2, Ad26.COV2.S, or NVX-CoV2373 COVID-19 vaccines at 6 months postvaccination (T5). Boolean analysis was carried out to define the functional profiles and the analysis included GzB, IFNγ, IL-2, and/or TNFα gated on CD3⁺CD8⁺ cells expressing CD69 (See **Figure 5**). Each Cytokine⁺ profile combination was considered positive with >0.005% and a SI>2 for CD8⁺ T cells. The dotted line indicates the limit of quantification (LOQ). The bars show the Geometric mean and geometric SD of the spike-specific CD8⁺ T cells CD69⁺.

T2, 15 ± 3 days; T3, 42 ± 7 days; T4, 108 ± 9 days; T5, 185 ± 8 days.

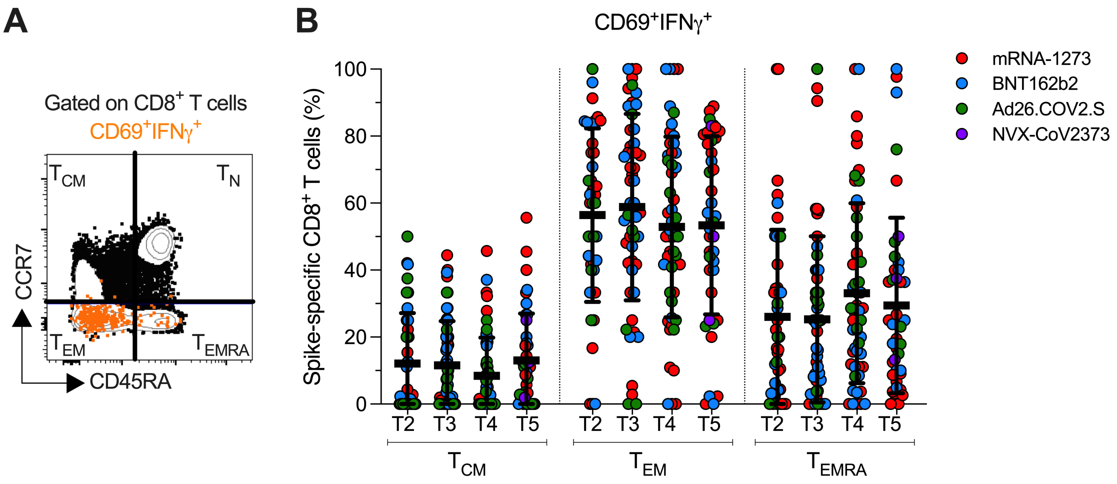


**Figure S10. Memory phenotype evaluated on spike-specific CD8⁺ T cells expressing CD69⁺ and producing IFNγ in subjects vaccinated with the mRNA-1273, BNT162b2, Ad26.COV2.S, or NVX-CoV2373 COVID-19 vaccines.**

**(A)** Representative gating strategy of the memory subsets evaluated on $CD8^+$ T cells expressing $CD69^+$ and producing IFNγ. The memory subsets were defined based on the expression of CCR7 and CD45RA: central memory ($T_{CM}$, $CCR7^+CD45RA^-$), effector memory ($T_{EM}$, CCR7-$CD45RA^-$), and terminally differentiated effector cells ($T_{EMRA}$, CCR7-$CD45RA^+$).

**(B)** Longitudinal analysis of the memory subsets evaluated on spike-specific $CD8^+$ T cells expressing $CD69^+$ and producing IFNγ in subjects vaccinated with the mRNA-1273, BNT162b2, Ad26.COV2.S, or NVX-CoV2373 COVID-19 vaccines.

T2, 15 ± 3 days; T3, 42 ± 7 days; T4, 108 ± 9 days; T5, 185 ± 8 days.

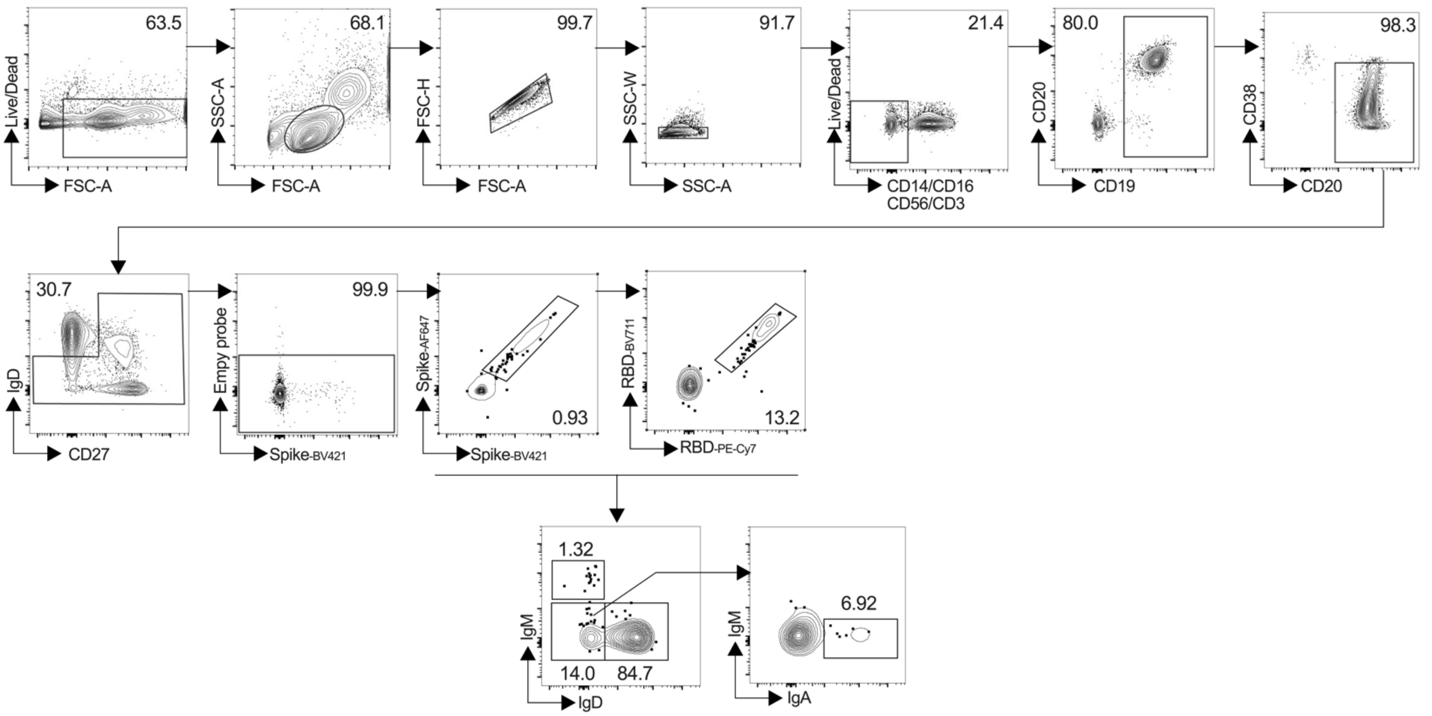


**Figure S11. Flow cytometry gating strategy for identification of spike and RBD-binding MBCs.**

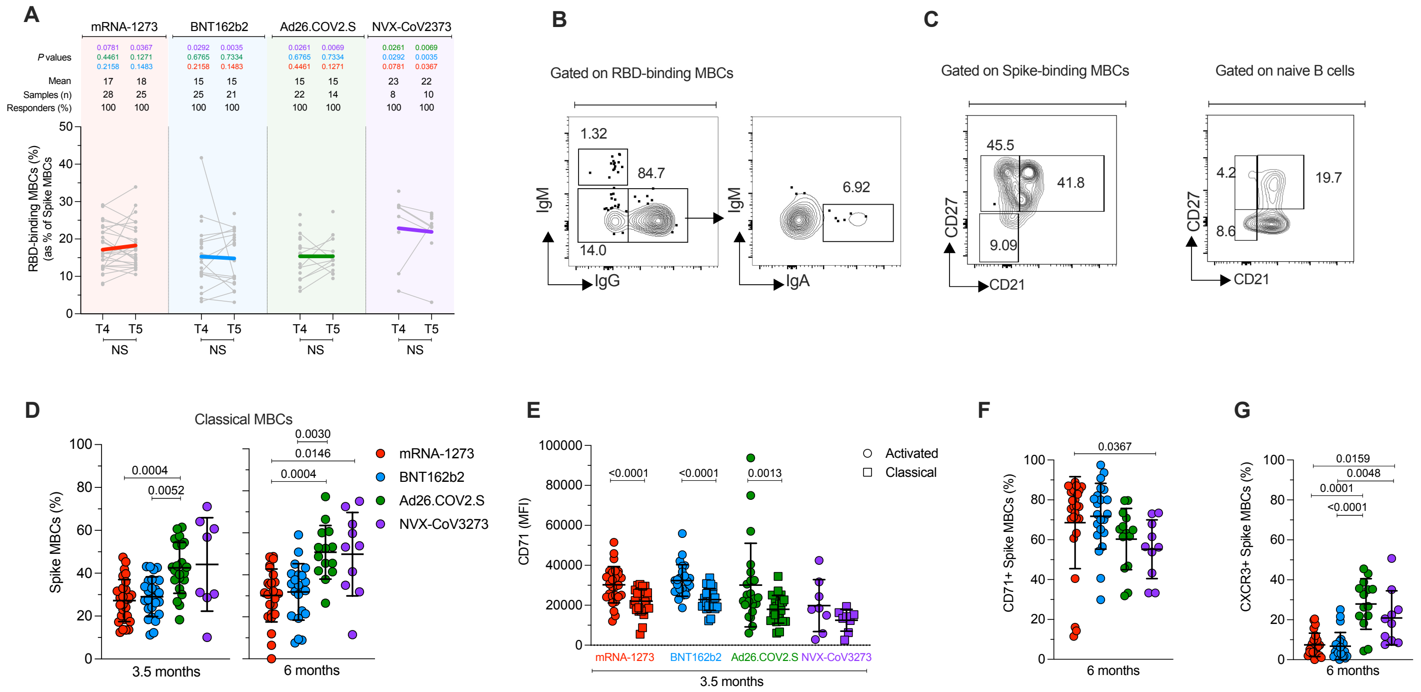

**Figure S12. Phenotypic characterization of SARS-CoV-2-specific MBCs induced by COVID-19 vaccine platforms.**
**(A)** Frequency of RBD MBCs gated from spike MBCs. P values on the top show the differences between each time point in the different vaccines. Bottom bars show T4 to T5 statistics.
**(B)** Representative gating strategy of Ig isotypes gated on RBD-binding MBCs.
**(C)** Representative gating strategy of spike MBCs with a classical ($CD21^+CD27^+$) and activated ($CD21^-CD27^+$) phenotype. Control gating on naïve B cells is shown for comparison.
**(D)** Frequency of classical spike MBCs ($CD21^+CD27^+$) at 3.5 (left panel) and 6 (right panel) months.
**(E)** CD71 expression by activated and classical spike-binding MBCs, at 3.5 months.
**(F-G)** Frequency of (F) $CD71^+$ and (G) $CXCR3^+$ spike-binding MBCs, at 6 months.
Data were analyzed for statistical significance using the Mann-Whitney test [(A), (E)], and Kruskal-Wallis (KW) test and Dunn's post-test for multiple comparisons [(D), (F), (G)]. NS, non-significant, MFI, median fluorescence intensity.

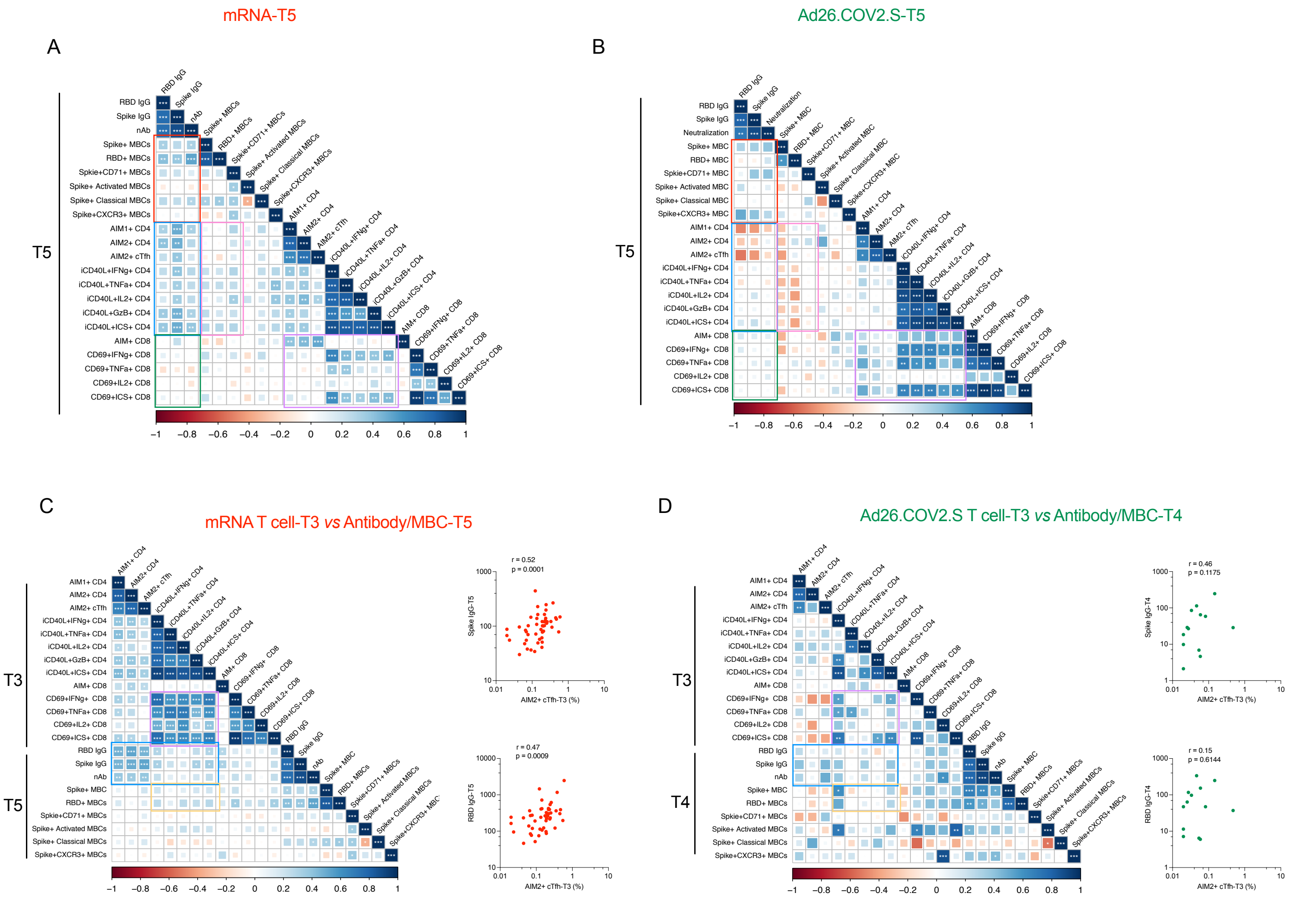


**Figure S13. Vaccine-specific correlation analyses.**

**(A-B)** Correlation matrix of T5 (6-month) samples, plotted as mRNA (mRNA-1273 and BNT162b2) and Ad26.COV2.S COVID-19 vaccines. The red rectangle indicated the association between antibody and memory B cells; the blue rectangle indicated the association between antibody and CD4+ T cell; the green rectangle indicated the association between antibody and CD8+ T cell; the pink rectangle indicated the association between CD4 T cells and memory B cell; the purple rectangle indicated the association between CD4 T cells and CD8 T cells. Spearman rank-order correlation values (r) are shown from red (-1.0) to blue (1.0); r values are indicated by color and square size. p values are indicated by white asterisks as * p <0.05, **p <0.01, *** p <0.001.

**(C-D)** Correlation matrix of CD4+ and CD8+ T cell data from the early time point with memory B cell and antibody data from the late timepoint. The blue rectangle indicates the association between CD4+ T cell and antibody; the orange rectangle indicated the association between CD4 T cells and memory B cell. Spearman rank-order correlation values (r) are shown from red (-1.0) to blue (1.0); r values are indicated by color and square size. P values are indicated by white asterisks as * p <0.05, **p <0.01, *** p <0.001. The T4 MBC and antibody data were preferred for Ad26.COV2.S due to fewer T5 paired samples.

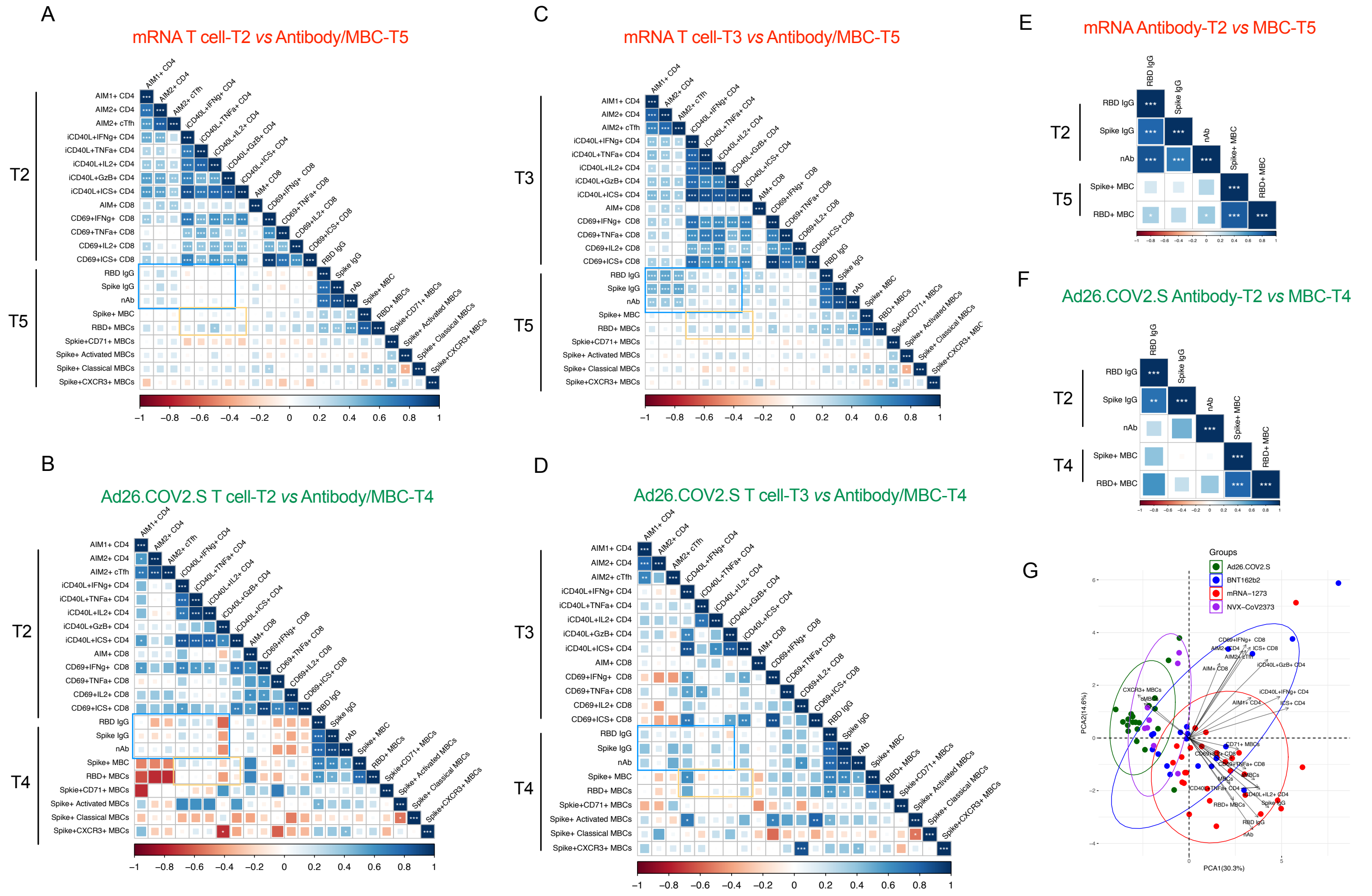

**Figure S14. Vaccine-specific correlation analyses.**

**(A-B)** Correlation matrix of CD4+ and CD8+ T cell data from the T2 time point with memory B cell and antibody data from the late timepoint. The blue rectangle indicates the association between CD4+ T cell and antibody; the orange rectangle indicated the association between CD4 T cells and memory B cell. Spearman rank-order correlation values (r) are shown from red (-1.0) to blue (1.0); r values are indicated by color and square size. P values are indicated by white asterisks as * p <0.05, **p <0.01, *** p <0.001. The T4 MBC and antibody data were preferred for Ad26.COV2.S due to fewer T5 paired samples.

**(C-D)** Correlation matrix of CD4+ and CD8+ T cell data from the T3 time point with memory B cell and antibody data from the late timepoint. The blue rectangle indicates the association between CD4+ T cell and antibody; the orange rectangle indicated the association between CD4 T cells and memory B cell. Spearman rank-order correlation values (r) are shown from red (-1.0) to blue (1.0); r values are indicated by color and square size. P values are indicated by white asterisks as * p <0.05, **p <0.01, *** p <0.001. The T4 MBC and antibody data were preferred for Ad26.COV2.S due to fewer T5 paired samples.

**(E-F)** Correlation matrix of antibody data from the T2 time point with memory B cell data from the late timepoint. Spearman rank-order correlation values (r) are shown from red (-1.0) to blue (1.0); r values are indicated by color and square size. P values are indicated by white asterisks as * p <0.05, **p <0.01, *** p <0.001. The T4 MBC data was preferred for Ad26.COV2.S due to fewer T5 paired samples.

**(G)** Principal component analysis (PCA) representation of mRNA-1273 (n=28), BNT162b2 (n=19), Ad26.COV2.S (n=20), and NVX-Cov-2373 (n=8) on the basis of all parameters obtained 3.5-month post-vaccination. Only paired subjects were used for the PCA analysis. Arrows indicated the prominent immunological distinguishing features. Ellipse represented the clustering of each vaccine. Red indicated mRANA-1273, blue indicated BNT162b2, and green indicated Ad26.COV2.S. MBCs indicates spike-specific memory B cell, cMBCs indicates spike-specific classical memory B cell, aMBCs indicates spike-specific activated memory B cell, AIM1+ indicates OX40+CD137+, AIM2+ indicates OX40+CD40L+, nAb indicates neutralization antibody.
